# Neurons of the inferior olive respond to broad classes of sensory input while subject to homeostatic control

**DOI:** 10.1101/379149

**Authors:** Chiheng Ju, Laurens W.J. Bosman, Tycho M. Hoogland, Arthiha Velauthapillai, Pavithra Murugesan, Pascal Warnaar, Romano M. van Genderen, Mario Negrello, Chris I. De Zeeuw

## Abstract

Cerebellar Purkinje cells integrate sensory information with motor efference copies to adapt movements to behavioural and environmental requirements. They produce complex spikes that are triggered by the activity of climbing fibres originating in neurons of the inferior olive. These complex spikes can shape the onset, amplitude and direction of movements as well as the adaptation of such movements to sensory feedback. Clusters of nearby inferior olive neurons project to parasagittally aligned stripes of Purkinje cells, referred to as “microzones”. It is currently unclear to what extent individual Purkinje cells within a single microzone integrate climbing fibre inputs from multiple sources of different sensory origins, and to what extent sensory-evoked climbing fibre responses depend on the strength and recent history of activation. Here we imaged complex spike responses in cerebellar lobule crus 1 to various types of sensory stimulation in awake mice. We find that different sensory modalities and receptive fields have a mild, but consistent, tendency to converge on individual Purkinje cells. Purkinje cells encoding the same stimulus show increased events with coherent complex spike firing and tend to lie close together. Moreover, whereas complex spike firing is only mildly affected by variations in stimulus strength, it strongly depends on the recent history of climbing fibre activity. Our data point towards a mechanism in the olivo-cerebellar system that regulates complex spike firing during mono- or multisensory stimulation around a relatively low set-point, highlighting an integrative coding scheme of complex spike firing under homeostatic control.

## Introduction

The olivo-cerebellar system is paramount for sensorimotor integration during motor behaviour. The climbing fibres that originate in the inferior olive and cause complex spike firing in cerebellar Purkinje cells encode both unexpected and expected sensory events and affect initiation, execution as well as adaptation of movements (Albus, 1971; Welsh *et al*., 1995; Kitazawa *et al*., 1998; Gibson *et al*., 2004; Bosman *et al*., 2010; Yang & Lisberger, 2014; Ohmae & Medina, 2015; Ten Brinke *et al*., 2015; Streng *et al*., 2017; Apps *et al*., 2018; Herzfeld *et al*., 2018; Junker *et al*., 2018). Complex spike firing frequency is typically sustained around 1 Hz and not substantially affected by the behavioural state, although short-lived increases or decreases in firing occur (Bloedel & Ebner, 1984; Mukamel *et al*., 2009; Bosman *et al*., 2010; Rahmati *et al*., 2014; Zhou *et al*., 2014; Hoogland *et al*., 2015; Negrello *et al*., 2018). To date, the paradox between the persistence of complex spike firing and the behavioural relevance of individual complex spikes is still largely unresolved.

Whereas the afferents of Purkinje cells, including not only the climbing fibres but also the parallel fibres and axons of interneurons, all diverge, their efferents strongly converge upon the cerebellar nuclei, ultimately integrating many different inputs from the brain (Harvey & Napper, 1991; Sugihara *et al*., 2001; Person & Raman, 2011). The parallel fibres are oriented in a transverse direction along the lobular axes, while the climbing fibres and axons of the interneurons are running perpendicularly to them in line with the sagittal orientation of the dendritic trees of Purkinje cells (Andersen *et al*., 1964; Szentágothai, 1965; Sugihara *et al*., 1999; Sullivan *et al*., 2005; Gao *et al*., 2006; Sugihara *et al*., 2009; Ruigrok, 2011; Cerminara *et al*., 2015; Apps *et al*., 2018). Interestingly, the intrinsic biochemical nature as well as the electrophysiological profile of individual Purkinje cells follows the organization of the climbing fibres so that Purkinje cells located in the same sagittal module receive climbing fibre input from the same olivary subnucleus and have similar identity properties, setting them apart from Purkinje cells in neighbouring modules (Xiao *et al*., 2014; Zhou *et al*., 2014; Cerminara *et al*., 2015; De Zeeuw & Ten Brinke, 2015; Tsutsumi *et al*., 2015; Suvrathan *et al*., 2016). Accordingly, Purkinje cell responses following electrical stimulation of major nerves of cat limbs largely adhere to the parasagittal organization of the climbing fibre zones (Oscarsson, 1969; Groenewegen *et al*., 1979), which in turn can be further differentiated into smaller microzones based upon their response pattern to tactile stimulation of a particular spot on the body (Ekerot *et al*., 1991; Apps & Garwicz, 2005; Ozden *et al*., 2009; De Zeeuw *et al*., 2011). Possibly, differential sensory maps even occur at the submicrozonal and individual Purkinje cell level, but this has to our knowledge not been investigated yet. In particular, it remains unclear to what extent different types of sensory input can drive complex spikes within the same individual Purkinje cells and/or their direct neighbours and how the strength as well as the history of these inputs influences the distribution of climbing fibre activity.

Here, we studied the impact of minimal stimuli of distinct sensory modalities on complex spike firing of Purkinje cells in crus 1 of awake mice using *in vivo* two-photon Ca^2+^ imaging with Cal-520 (Tada *et al*., 2014). This fluorescent sensor has been reported to be a more accurate reporter of fast spikes than commonly used genetically encoded sensors such as GCaMP6f (Lock *et al*., 2015). We found that different sensory streams appear to converge on individual inferior olivary neurons and thereby Purkinje cells, that sensory stimulation primarily affects the timing of the complex spikes rather than their rate, that the strength of complex spike responses varied seamlessly from non-responsive to highly responsive, that a recent history of high activity leads to a future of low activity, and that Purkinje cells that respond to the same stimulus tend to be located in each other’s vicinity and display increased levels of simultaneous firing. Together, our data indicate that subtle and local sensory inputs can recruit mosaic ensembles of Purkinje cells, employing population coding in a spatially and temporally dynamic way that is in line with homeostatic control.

## Methods

### Ethical approval

All experimental procedures involving animals were in agreement with Dutch and European legislation and guidelines as well as with the ethical principles of The Journal of Physiology. The experiments were approved in advance by an independent ethical committee (DEC Consult, Soest, The Netherlands) as required by Dutch law and filed with approval numbers EMC2656, EMC3001 and EMC3168. Experiments were performed in compliance with the guidelines of the Animal Welfare Board of the Erasmus MC.

### Animals and surgery

Mice were group housed until the day of the experiment and kept under a regime with 12 h light and 12 h dark with food and water available *ad libitum*. The mice had not been used for other experiments prior to the ones described here.

For the experiments performed in awake mice, we recorded from in total 66 field of views located in cerebellar lobule crus 1 of 29 male C57BL/6J mice of 4-12 weeks of age (Charles Rivers, Leiden, The Netherlands). Prior to surgery, mice were anaesthetized using isoflurane (initial concentration: 4% V/V in O_2_, maintenance concentration: ca. 2% V/V) and received Carprofen (Rimadyl, 5 mg/ml subcutaneously) to reduce post-surgical pain. Before the start of the surgery, the depth of anaesthesia was verified by the absence of a reaction to an ear pinch. To prevent dehydration, mice received 1 ml of saline s.c. injection before the surgeries commenced. Eyes were protected using eye ointment (Duratears, Alcon, Fort Worth, TX, USA). Body temperature was maintained using a heating pad in combination with a rectal thermometer. During surgery, we attached a metal head plate to the skull with dental cement (Superbond C&B, Sun Medical Co., Moriyama City, Japan) and made a craniotomy with a diameter of approximately 2 mm centred on the medial part of crus 1 ipsilateral to the side of somatosensory stimulation. The dura mater was preserved and the surface of the cerebellar cortex was cleaned with extracellular solution composed of (in mM) 150 NaCl, 2.5 KCl, 2 CaCl_2_, 1 MgCl_2_ and 10 HEPES (pH 7.4, adjusted with NaOH). After surgery, the mice were allowed to recover from anaesthesia for at least 30 minutes. Subsequently, the mice were head-fixed in the recording setup and they received a bolus-loading of the Ca^2+^ indicator Cal-520 (0.2 mM; AAT Bioquest, Sunnyvale, CA) (Tada *et al*., 2014). Cal-520 was used as it is currently the best available green Ca^2+^ dye, outperforming genetically encoded indicators such as GCaMP6f (Lock *et al*., 2015). The dye was first dissolved with 10% w/V Pluronic F-127 in DMSO (Invitrogen, Thermo Fisher Scientific, Waltham, MA, USA) and then diluted 20 times in the extracellular solution. The dye solution was pressure injected into the molecular layer (50–80 μm below the surface) at 0.35 bar for 5 min. Finally, the brain surface was covered with 2% agarose dissolved in saline (0.9% NaCl) in order to reduce motion artefacts and prevent dehydration.

For the experiments on single-whisker stimulation we made recordings under anaesthesia on 17 male C57BL/6J mice of 4-12 weeks of age. The procedure was largely the same as described above, but instead of isoflurane we used ketamine/xylazine as anaesthetic (i.p. injection via butterfly needle, initial dose: 100 mg/kg and 10 mg/kg, respectively; maintenance dose: approximately 60 mg/kg/h and 3 mg/kg/h, respectively). The mice remained under anaesthesia until the end of the recording. A subset of these experiments was performed with 0.2 mM Oregon Green BAPTA-1 AM dye (Invitrogen) as this dye has been more widely used than Cal-520 (e.g., Stosiek *et al*., 2003; Ozden *et al*., 2009; Schultz *et al*., 2009; Hoogland *et al*., 2015). Oregon Green BAPTA-1 AM was dissolved and applied in the same way as Cal-520. We found that Cal-520 had a superior signal-to-noise ratio under our experimental conditions. As a consequence, the observed event rate was lower in the experiments using OGB-1 (OGB-1: 0.45 ± 0.26 Hz; *n* = 172 cells; Cal-520: 0.72 ± 0.40 Hz; *n* = 43 cells; median ± IQR; *U* = 1719.0, *p* < 0.001, Mann-Whitney test). The observed frequency range using Cal-520 was comparable to that found using *in vivo* single-unit recordings under ketamine/xylazine anaesthesia: 0.6 ± 0.1 Hz (Bosman *et al*., 2010). Despite an underestimation of the complex spike rate using OGB-1, we found that the ratios of Purkinje cells that responded to single whisker stimulation were similar for both dyes (OGB-1: 89 out of 373 cells (24%); Cal-520: 35 out of 152 cells (23%); *p* = 0.910; Fisher’s exact test). For this reason, we combined the data from both dyes. For the analysis presented in Fig. 11, we included only those cells that could be recorded during all stimulus conditions. All experiments using awake data were obtained with Cal-520.

At the end of each experiment, the mice were killed by cervical dislocation under isoflurane or ketamine/xylazine anaesthesia after which the brain was removed and the location of the dye injection in crus 1 was verified by epi-fluorescent imaging. The whole procedure, from initial anaesthesia to cervical dislocation, typically lasted around 6 to 8 h.

### In vivo *two-photon Ca2+ imaging*

Starting at least 30 min after dye injection, *in vivo* two-photon Ca^2+^ imaging was performed of the molecular layer of crus 1 using a setup consisting of a Ti:Sapphire laser (Chameleon Ultra, Coherent, Santa Clara, CA, USA), a TriM Scope II system (LaVisionBioTec, Bielefeld, Germany) mounted on a BX51 microscope with a 20x 1.0 NA water immersion objective (Olympus, Tokyo, Japan) and GaAsP photomultiplier detectors (Hamamatsu, Iwata City, Japan). A typical recording sampled a field of view of 40 × 200 µm with a frame rate of approximately 25 Hz.

In a subset of experiments (Fig. 12A-C), larger field-of-views were obtained using a two-photon setup from Neurolabware (Los Angeles, CA, USA). Imaging occurred through a 16x (0.8 NA) objective (Nikon, Tokyo, Japan) in combination with a Chameleon Ultra II laser (Coherent) tuned to 920 nm at a frame rate of 15 Hz.

### Sensory stimulation

Cutaneous stimuli were delivered to four defined regions on the left side of the face, ipsilateral to side of the craniotomy. These regions were the whisker pad, the cheek posterior to the whisker pad, the upper lip and the lower lip. Stimuli were applied using a Von Frey filament (Touch Test Sensory Evaluator 2.83, Stoelting Co., IL, USA) attached to a piezo linear drive (M-663, Physik Instrumente, Karlsruhe, Germany). Prior to the set of experiments described here, we tested a series of 8 Von Frey filaments with a stiffness range from 0.02 g to 1.4 g in awake head-fixed mice to select the optimal force for these experiments. We selected the 0.07 g (0.686 mN) filament because this filament induced a mild reaction in the mouse, but no signs of a nociceptive response (cf. Chaplan *et al*. (1994)). The touch time was fixed at 100 ms. As a control, we moved the stimulator without touching the face (“sound only” condition). Visual stimuli were delivered as 10 ms pulses using a 460 nm LED (L-7104QBC-D, Kingbright, CA, USA). The stimulation frequency was fixed at 1 Hz and the different stimuli were applied in a random order. Video recordings of the eye, made under infrared illumination, revealed that the mice did not make eye movements in response to the LED flash, but they did show reflexive pupil constriction (*data not shown*).

Single whiskers were stimulated using a piezo linear drive (M-663, Physik Instrumente) while using a deflection of 6°. It took the stimulator approximately 30 ms to reach this position. At the extreme position, the stimulator was paused for 150 ms before returning to the neutral position. The stimulator was designed to minimize contact with other whiskers. Each stimulation experiment consisted of five sessions in random order. During each block one of the ipsilateral whiskers B2, C1, C2, C3 and D2 was stimulated. Each session consisted of 150 stimuli at 2 or 3 Hz.

### Complex spike detection

Image analysis was performed offline using custom made software as described and validated previously (Ozden *et al*., 2008; Ozden *et al*., 2012; De Gruijl *et al*., 2014). In short, independent component analysis was applied to the image stack to discover masks describing the locations of individual Purkinje cell dendrites (e.g., see Fig. 2A). For each field of view, the mask was generated only once, so that the same Purkinje cells were analysed for subsequent recordings of different stimulus conditions enabling paired comparisons. Experiments during which spatial drift occurred were discarded from subsequent analysis. The fluorescence values of all pixels in each mask were averaged per frame. An 8% rolling baseline from a time window of 0.5 ms was subtracted from the average fluorescence per mask (Ozden *et al*. 2012), after which Ca^2+^ transient events were detected using template matching.

### Statistical analysis

In general, we first tested whether parameter values were distributed normally using one-sample Kolmogorov-Smirnov or Shapiro-Wilk tests. If not, non-parametric tests were applied. When multiple tests were used, Benjamini-Hochberg correction for multiple comparisons was applied. For each experiment, stimuli were given in a random sequence. Identification of Purkinje cell dendrites and event extraction were performed by a researcher who was blind to the type of stimulus. For the experiments characterizing response characteristics of individual Purkinje cells, we compared the fraction of responsive Purkinje cells, the response latency and the response peak. After extracting the Ca^2+^ transient event times, peri-stimulus time histograms (PSTHs) were constructed using the inter-frame time (approx. 40 ms) as bin size. Stacked line plots of Purkinje cell PSTHs were sorted by weak to strong peak responses. The data were normalized such that the top represents the average responses of all Purkinje cells. Statistical significance of responses occurring within 200 ms from stimulus onset was evaluated using a threshold of the mean + 3 s.d. of the firing rate during the 500 ms prior to stimulus onset (300 ms for the single whisker stimulation experiments, as they were performed with a higher stimulation frequency).

To calculate whether two or more inputs converged on single Purkinje cells we used a bootstrap method. First, we determined the fraction of responsive cells for each parameter and compared these to a randomly generated number between 0 and 1 as taken from a uniform distribution. If for each input the randomly generated numbers were lower than the measured fractions, we considered them as responsive to all stimuli. This procedure was repeated 10,000 times and the average and standard deviation were derived and used to calculate the *Z* score of the experimental data. All bootstrap procedures were performed using custom-written code in LabVIEW (National Instruments, Austin, TX, USA).

A direct comparison between the full and partial correlations of the peak responses to tactile stimuli was performed in MATLAB (MathWorks, Natick, MA, USA) (Fig. 8A-B). Principal component analysis of the same dataset was also performed in MATLAB and compared to that of a bootstrapped dataset (Fig. 8C). The latter was obtained by shuffling the response peaks per stimulus condition 500 times. After each shuffle, a principal component analysis was performed of which the average (± 3 s.d.) was calculated and plotted. The analyses described in Fig. 8A-C were performed on those 188 Purkinje cells that received all four tactile stimuli.

The distributions of pairs of Purkinje cells, either both responsive to a given stimulus (“responsive pairs”) or one cell being responsive and the other not (“heterogeneous pairs”), were tested using two-dimensional Kolmogorov-Smirnov tests performed in MATLAB. Aggregate PSTHs (Fig. 13) were constructed and evaluated as described before (Romano *et al*., 2018). Briefly, we calculated per individual frame the number of simultaneously occurring events and colour coded these in a PSTH combining data from all dendrites in a field of view. Based upon the total number of complex spikes and dendrites per recording, we calculated the expected number of simultaneous complex spikes per individual frame using a Poisson distribution. The actual number of simultaneous complex spikes was compared to this calculated distribution and a *p* value was derived for each number based upon the Poisson distribution (using custom-written software in MATLAB and LabVIEW). Unless mentioned otherwise, correlation analysis was performed in SigmaPlot (Systat Software, San Jose, CA, USA) and the other statistical tests were performed using SPSS (IBM, Armonk, NY, USA).

## Results

### Climbing fibre responses to tactile, auditory and/or visual input

Little is known about the distribution and convergence of different types of climbing fibre-mediated sensory input at the level of individual, nearby Purkinje cells. Here, we performed two-photon Ca^2+^ imaging of Purkinje cells of awake mice to record complex spikes responses to various types of sensory stimulation related to the face of the mice. The recordings were made in crus 1 as this lobule is known to receive sensory input from the oro-facial region, in particular from the mystacial vibrissae (Fig. 1A) (Shambes *et al*., 1978; De Zeeuw *et al*., 1990; Yatim *et al*., 1996; Apps & Hawkes, 2009; Bosman *et al*., 2011; De Gruijl *et al*., 2013; Kubo *et al*., 2018; Romano *et al*., 2018). The configurations and positions of individual Purkinje cell dendrites were detected using independent component analysis (Fig. 1B). Ca^2+^ transients were isolated per dendrite by a template-matching procedure that reliably detects complex spikes (Ozden *et al*., 2008; De Gruijl *et al*., 2014; Najafi *et al*., 2014) (Fig. 1C-E).

**Figure 1.**
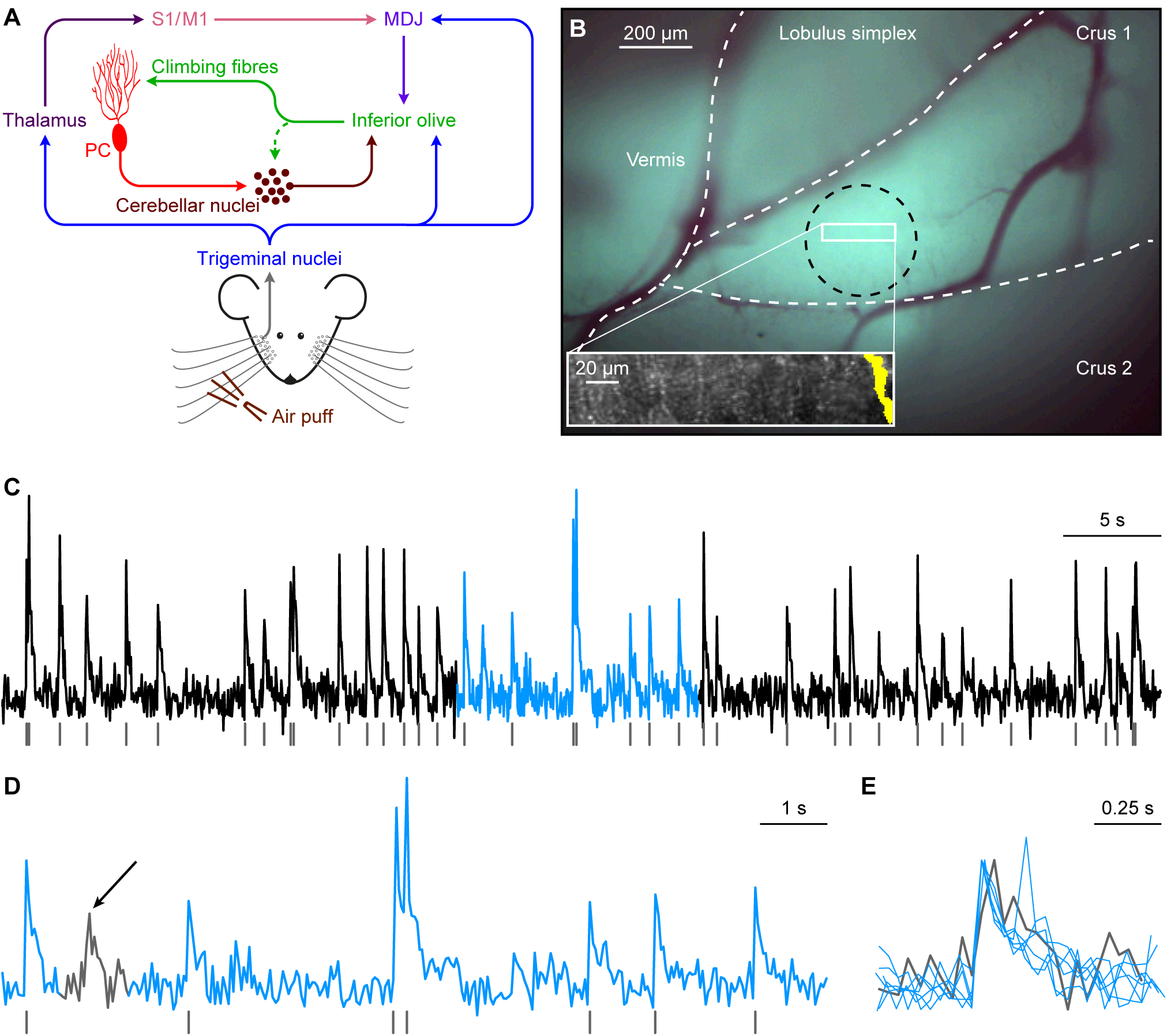
Sensory pathways carrying facial input to the cerebellar cortex. **(A)** Scheme of the main routes conveying facial tactile input via the climbing fibre pathway to cerebellar Purkinje cells (PC). Climbing fibres, which cause complex spike firing in Purkinje cells, exclusively originate from the inferior olive. The inferior olive, in turn, is directly innervated by neurons from the trigeminal nuclei as well as indirectly via thalamo-cortical pathways that project to the inferior olive mainly via the nuclei of the mesodiencephalic junction (MDJ). The MDJ itself also receives direct input from the trigeminal nuclei. See the main text for references. **(B)** *In vivo* two-photon Ca^2+^ imaging was performed to characterize Purkinje cell complex spike responses to sensory stimulation in the medial part of crus 1. Purkinje cells were detected using independent component analysis and the position of a Purkinje cell dendrite (yellow area on the right) within a field of view is shown in the inset. At the end of each recording session, the brain was removed and the location of the dye injection in medial crus 1 was confirmed through *ex vivo* epifluorescent imaging (black circle). The white rectangle indicates the approximate recording location. **(C)** Complex spikes that were triggered by climbing fibre activity were retrieved from fluorescent traces of individual Purkinje cells. A representative trace obtained from the Purkinje cell dendrite illustrated in **B** is shown together with the detected complex spikes (grey lines). The light blue episode is enlarged in **D**. Complex spikes were detected by the combination of a threshold and a template matching algorithm. Only events with a sharp rising phase were accepted as complex spikes. In the 60 s interval shown in **C**, there was one event with a slower rise time (see arrow in **D**), as indicated at a larger time scale in **E**. The events in **E** are scaled to peak.

**Figure 2.**
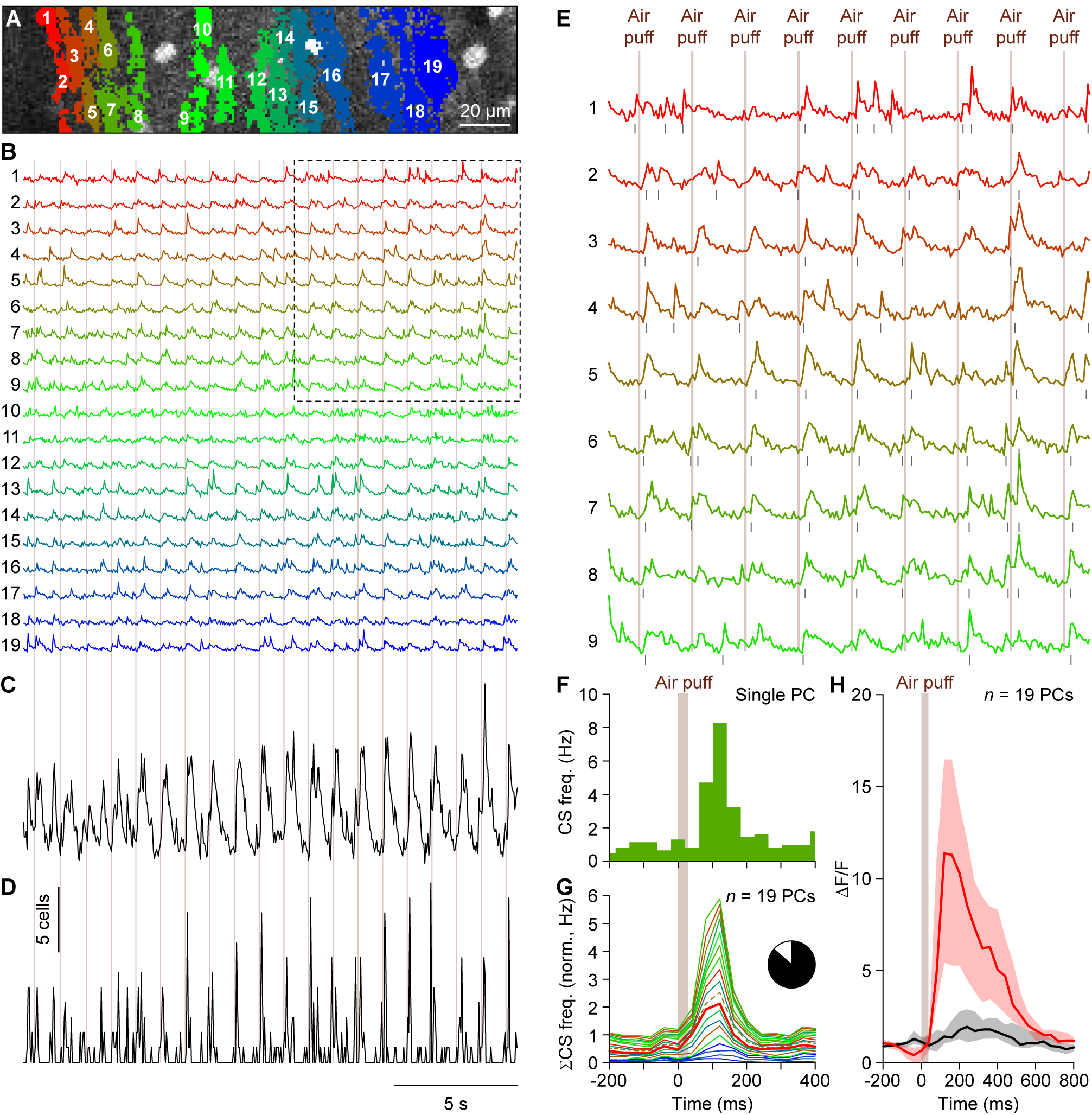
Whisker stimulation evokes complex spike responses in cerebellar crus 1. **(A)** Complex spikes elicit large increases in the Ca^2+^ concentration within Purkinje cell dendrites that can be resolved using *in vivo* two-photon microscopy in combination with a fluorescent Ca^2+^ indicator. An example of a field of view with 19 identified Purkinje cell dendrites located in the medial part of crus 1 is shown with each individual dendrite denoted by a number and a unique colour. This recording was made in an awake mouse. **(B)** The fluorescent traces of each of these dendrites show distinct Ca^2+^ transient events, which are to a large extent associated with air puff stimulation of the facial whiskers (times of stimulation indicated by the vertical lines). The boxed area is enlarged in **E**. **(C)** Summed fluorescence trace composed of all 19 individual traces emphasizing the participation of many Purkinje cells to the stimulus-triggered responses. **(D)** After complex spike extraction, a clear relation between stimulus and activity was observed as illustrated by summing, for each frame, the number of complex spikes observed over all dendrites. The vertical scale bar corresponds to the simultaneous activity of 5 Purkinje cells. **(E)** Enlargement of the boxed area in **B**. The air puffs were delivered once every second. **(F)** Peri-stimulus time histogram of a Purkinje cell dendrite (marked as number 7 in **A** and **B**) in response to air puff stimulation to the ipsilateral whiskers (154 trials). The bin size (40 ms) corresponds to the acquisition frame rate (25 Hz). **(G)** Normalized stacked line graph of the Purkinje cells in this field of view showing that every cell contributed to the overall response. The Purkinje cells are ranked by their maximal response and the data are normalized so that the top line reflects the average frequency per bin. Cell no. 5 (dashed line) had a relatively poor signal-to-noise ratio during later parts of the recording (see Fig. 3), but it had nevertheless a complex spike response profile that was indistinguishable from the other cells. The colours match those in the panels **A**-**B**. Inset: In total, 102 out of 117 cells analysed (87%) were responsive to whisker air puff stimulation (peak response exceeded average + 3 s.d. of pre-stimulus interval). **(H)** Median fluorescent traces of the trials with (red) and without (black) complex spikes fired during the first 200 ms after air puff onset. In the absence of complex spike firing, only a very small increase in fluorescence was observed, indicating that the majority of change in fluorescence was associated with complex spike firing. Note the longer time scale than in **F** and **G**. The lines indicate the medians and the shaded areas the inter-quartile ranges of the 19 Purkinje cells in this field of view.

We started our study of sensory responses by using air puff stimulation of the large facial whiskers, which is a relatively strong stimulus. In line with previous studies (Axelrad & Crepel, 1977; Brown & Bower, 2002; Bosman *et al*., 2010; Apps *et al*., 2018; Romano *et al*., 2018), we found that air puff stimulation to the whiskers evoked complex spike responses in many Purkinje cells (e.g., in 19 out of 19 Purkinje cells in the example illustrated in Fig. 2). In total, 102 out of the 117 (87%) cells analysed were considered to be responsive to such a stimulus, implying that the peak responses of these cells exceeded the threshold of the average + 3 s.d. of the pre-stimulus period.

Complex spikes are associated with a fast and large increase in intracellular Ca^2+^ in the dendrites of Purkinje cells, but other processes may also contribute to a rise in the intracellular Ca^2+^ concentration (Najafi *et al*., 2014; Roome & Kuhn, 2018). Indeed, in our data, we could observe changes in fluorescence that were not related to complex spikes (Fig. 2H). To exclude the possibility that we erroneously detected such non-complex spike events as complex spikes, we overlaid trials with and without complex spikes. This revealed that fluorescent events that were not classified as complex spikes typically had smaller amplitudes and slower kinetics than complex spikes, with the exception of a very noisy recording where non-complex spikes were much faster than complex spikes (Fig. 3A). This confirms the impression revealed by Fig. 1D-E that complex spikes can be properly detected based upon their waveform, in line with previous reports using the same detection algorithm (Ozden *et al*., 2008; De Gruijl *et al*., 2014; Najafi *et al*., 2014). To further characterise the rise times, we performed a set of experiments with a smaller field of view and a higher frame rate (100 Hz). These recordings revealed that virtually all events had a rise time of 1 or 2 frames (10 or 20 ms) (Fig. 3B-D).

**Figure 3.**
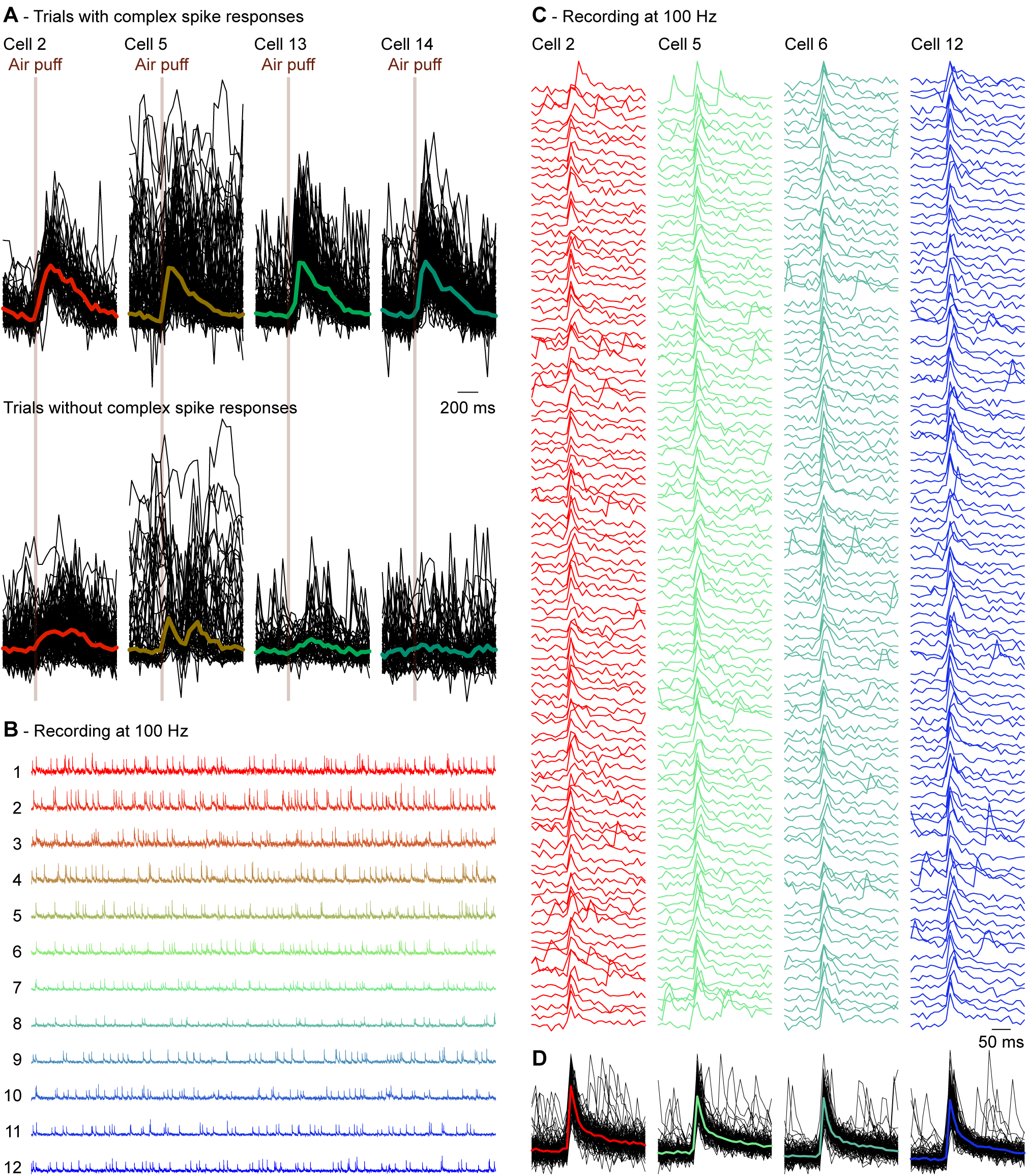
Complex spike detection. **(A)** Of the same experiment illustrated in Fig. 2, we randomly selected four Purkinje cells of which all trials with (top) and without (bottom) a complex spike within 200 ms of stimulus onset are shown. The signals are scaled to the maximum of the median complex spike responses. The fat coloured lines indicate the medians with the same colour code as in Fig. 2. Trials without a complex spike response can still show an increased fluorescence, but the kinetics of the non-complex spike response differed from those of the complex spike responses. Note that the raw trace of cell 5 had a period with increased noise levels, making it the least reliable cell of this recording. Nevertheless, cell 5 had a complex spike response profile that was very similar to those of the other cells (see dashed line in Fig. 2G). Cells (in other recordings) that had a worse signal-to-noise ratio than cell 5 of this recording were excluded from analysis. **(B)** To have a better characterization of the rise times, we made recordings with a smaller field of view, but a higher temporal resolution (100 Hz). Shown are 60 s traces of 12 simultaneously recorded Purkinje cells. **(C)** Of four randomly selected Purkinje cells, the first 100 events were plotted and superimposed **(D)**. The coloured lines in **D** indicate the medians. Virtually all events had a rise time of 10 or 20 ms (1 or 2 frames).

To address whether localized and more subtle stimuli recruited smaller groups of Purkinje cells, maybe even subsets of microzones, we subsequently applied gentle tactile stimulation at four facial locations: the whisker pad, the cheek posterior to the whisker pad, the upper lip and the lower lip (Fig. 4A). Although the precise somatotopy of climbing fibre projections to crus 1 is not known, it has been shown that the mossy fibre projections to crus 1 convey somatosensory input mainly from the mystacial vibrissae and the surrounding skin, while the somatosensory input from the lips predominantly targets crus 2 (Shambes *et al*., 1978). The stimuli were given using a Von Frey filament (target force = 0.686 mN) attached to a piezo actuator. The stimulus strength was carefully calibrated to avoid inducing responses from neighbouring skin areas or nociceptive responses (see Methods). The strength of the Purkinje cell responses to any of the four tactile stimuli (998 stimulus conditions in 282 Purkinje cells) had a skewed, but continuous distribution (Fig. 4B). Moreover, for individual stimulus locations such skewed distributions of response strengths were found as well, with the upper lip being the least sensitive of the four facial areas (Fig. 5). In other words, our data set contained a gradient of both non-responsive and strongly responsive Purkinje cells. A distinction between “responsive” and “non-responsive” Purkinje cells therefore involved a somewhat arbitrary distinction, prompting us to present most of the subsequent analyses using the entire dataset.

**Figure 4.**
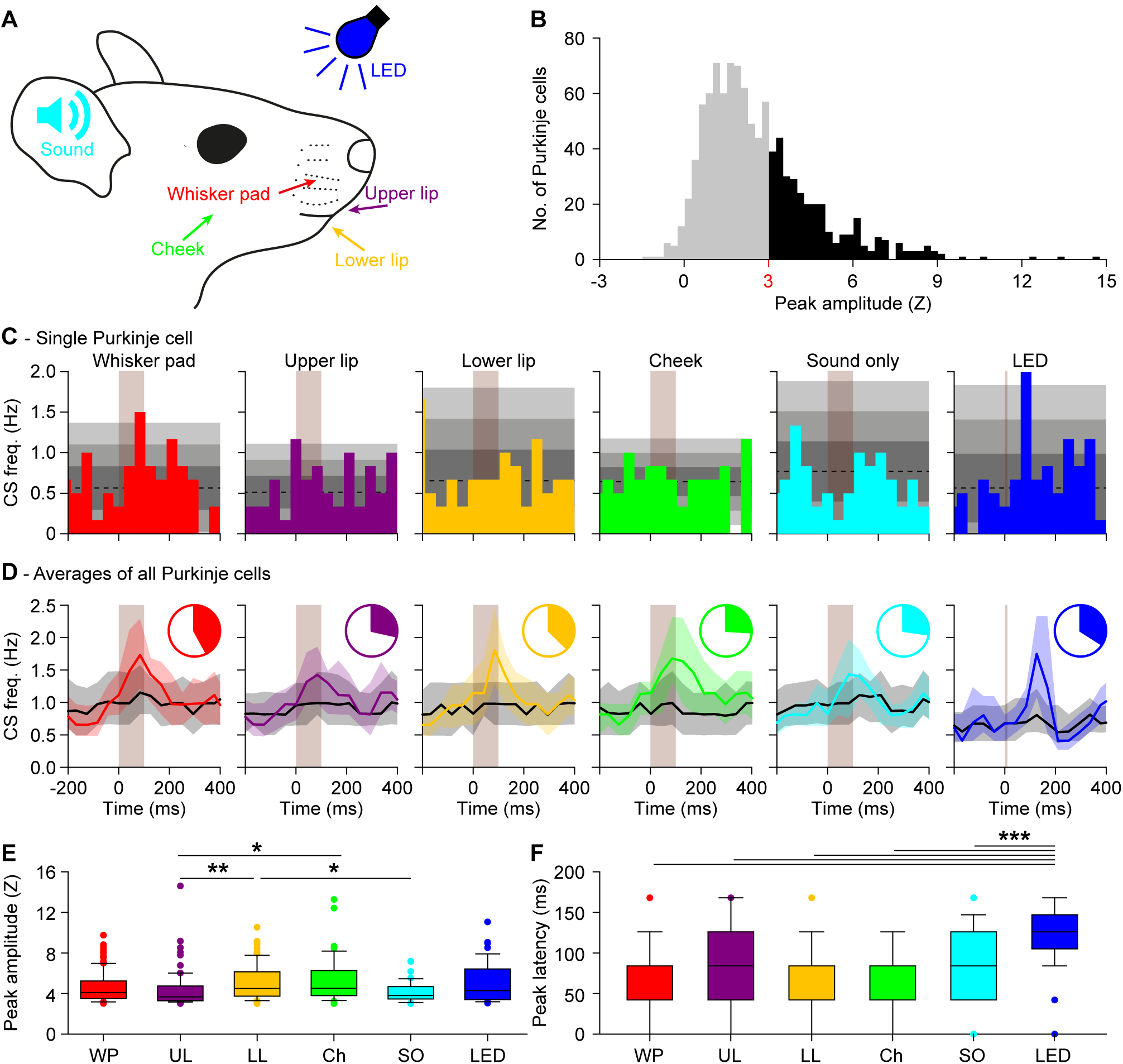
Purkinje cells in crus 1 respond to various types of sensory stimulation. **(A)** Tactile stimuli were presented to four facial regions in awake mice: the whisker pad (WP), the upper lip (UL), the lower lip (LL) and the cheek (Ch). Each stimulus was delivered with a piezo-actuator that made a muted, yet audible sound which was also delivered without touch (“sound only (SO)”). A blue light flash generated by an LED was used as visual stimulus. These experiments were performed in awake mice. **(B)** To avoid interference of adjacent areas, we applied gentle touches (0.686 mN). The complex spike response ratio was much reduced relative to the strong air puff stimulation to all ipsilateral whiskers illustrated in Fig. 2. A histogram of the peak responses (expressed as Z value) of all responses to either of the four tactile stimuli demonstrates that the response strength is a continuum, showing the lack of a clear separation between “responsive” and “non-responsive” Purkinje cells (998 stimulus conditions in 282 Purkinje cells). We considered Purkinje cells that showed a peak response above Z = 3 as “significantly responsive” (represented with black bars), but we provide most of the analyses also for the population as a whole (e.g., Fig. 5). **(C)** The peri-stimulus time histograms (PSTHs) of a representative Purkinje cell. The shades of grey indicate 1, 2 and 3 s.d. around the average. Each stimulus was repeated 154 times at 1 Hz. **(D)** For every stimulus condition, we averaged the PSTHs for all Purkinje cells that were significantly responsive to that particular stimulus (coloured lines; medians (inter-quartile range)). These were contrasted to the averaged PSTH of the other Purkinje cells (black lines). The pie charts represent the fraction of Purkinje cells significantly responsive to a particular stimulus. See also Table 1. **(E)** The peak responses of the significantly responding Purkinje cells were the lowest for sound only and for upper lip stimulation. * *p* < 0.05; ** *p* < 0.01 (post-hoc tests after Kruskal-Wallis test) **(F)** As expected for complex spike responses to weak stimulation, the latencies were relatively long and variable, but consistent across types of stimulation. Only visual stimulation (LED) had a remarkably longer latency time. *** *p* < 0.001 (post-hoc tests after Kruskal-Wallis test for LED vs. whisker pad, upper lip, lower lip and cheek and *p* < 0.05 compared to sound only)

**Figure 5.**
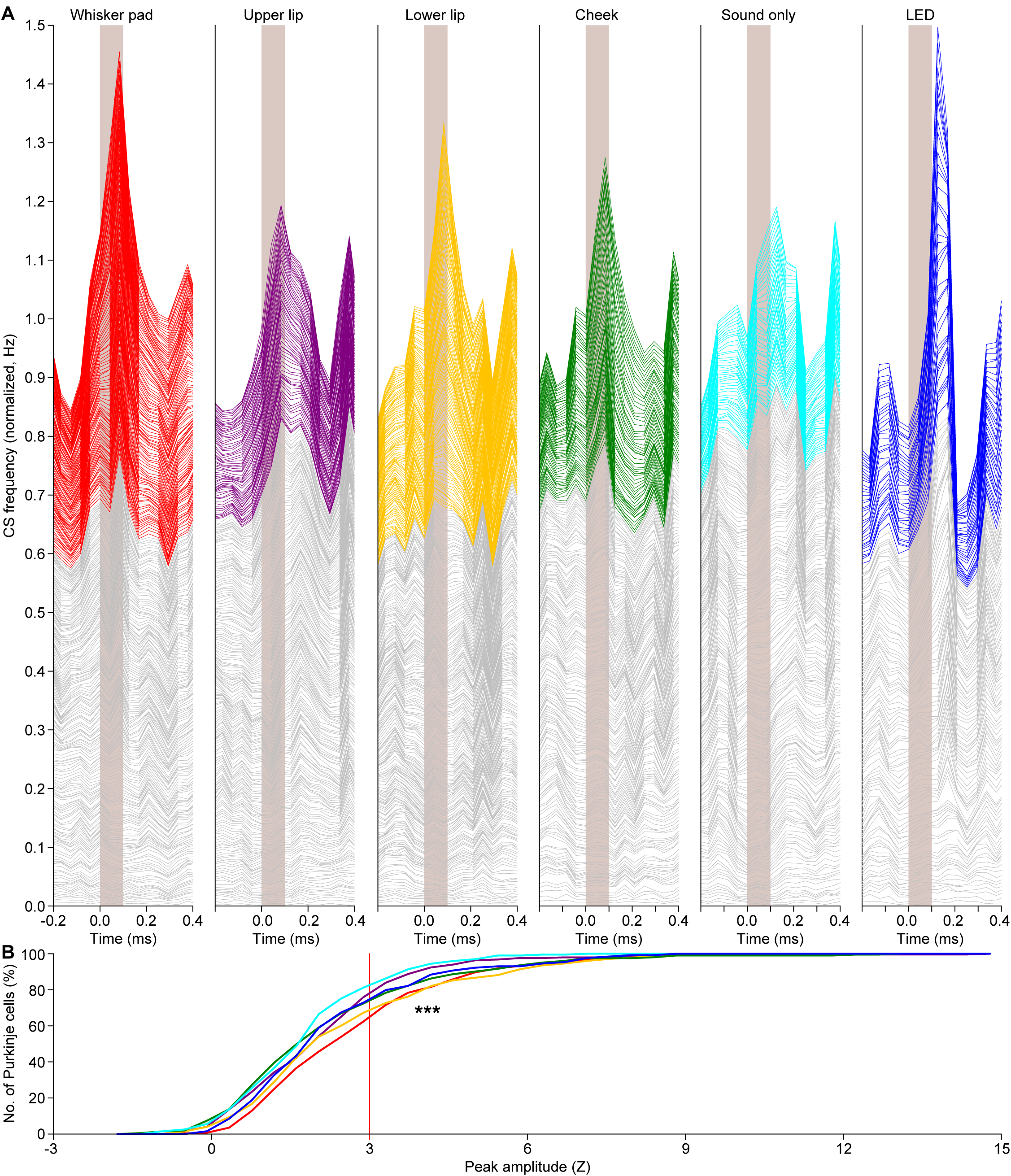
Variations in response strength. **(A)** Stacked line plots illustrating the peri-stimulus time histograms of all Purkinje cells recorded under either of the six indicated stimulus conditions. The cells are ordered based upon their peak responses (calculated as Z value) during the 200 ms interval following the stimulus onset, with the cell with the lowest response at the bottom of each graph. The grey lines indicate cells with a peak response deemed not significant (Z < 3) and the coloured lines indicate the significant responses (Z > 3). The graphs are normalized so that the upper line depicts the averge of all cells. As shown in Fig. 4B, one cannot discriminate between responsive and non-responsive cells in a black-and-white fashion. Instead, the cells form a continuum from not responsive at all to highly responsive. As this way of plotting relies on the numerical average, a skewed distribution can put emphasis on a relatively small group of cells. For this reason, the Purkinje cell responses are also compared using cumulative histograms **(B)** using the same colour scheme as in **A**. From this representation, it is confirmed that whisker pad and lower lip stimulation yield the strongest responses, while visual stimulation (LED) recruits a few cells with a relatively strong response, increasing the numerical average (see **A**). Here, we tested the complete distributions (*** *p* < 0.001; Kruskal-Wallis test). Pair-wise comparisons of all stimulus conditions are presented in Fig. 6.

Given that touches were delivered by a piezo-actuator that made a weak but audible sound, we also tested whether Purkinje cells responded to this sound in the absence of touch. This was the case, and, as expected, the “sound only” stimulus evoked the weakest responses of all stimuli (*p* < 0.001; Kruskal-Wallis test; Fig. 5B). For comparison, we also included a visual stimulation, consisting of a brief (10 ms) flash of a blue LED. This stimulus evoked responses with a similar strength as the whisker pad and lower lip stimulation, but with a latency that was remarkably long (Fig. 5A).

To facilitate a quantitative comparison between the stimulus conditions, we subsequently focused on the subset of obvious responses, defined as having a peak amplitude exceeding our threshold for significance set at the average + 3 s.d. of the pre-stimulus period. Among these “significantly responding” Purkinje cells, we found trends as in the entire population: the lower lip recruited the strongest responses, directly followed by the whisker pad and visual stimulation, while upper lip and sound only stimulations were less effective (Fig. 4C-E; Table 1). We also confirmed the remarkably long latency, typically more than 100 ms, for the visually evoked responses (*p* < 0.001 compared to the tactile stimuli; Kruskal Wallis test; Fig. 4F; Table 1).

**Table 1.**
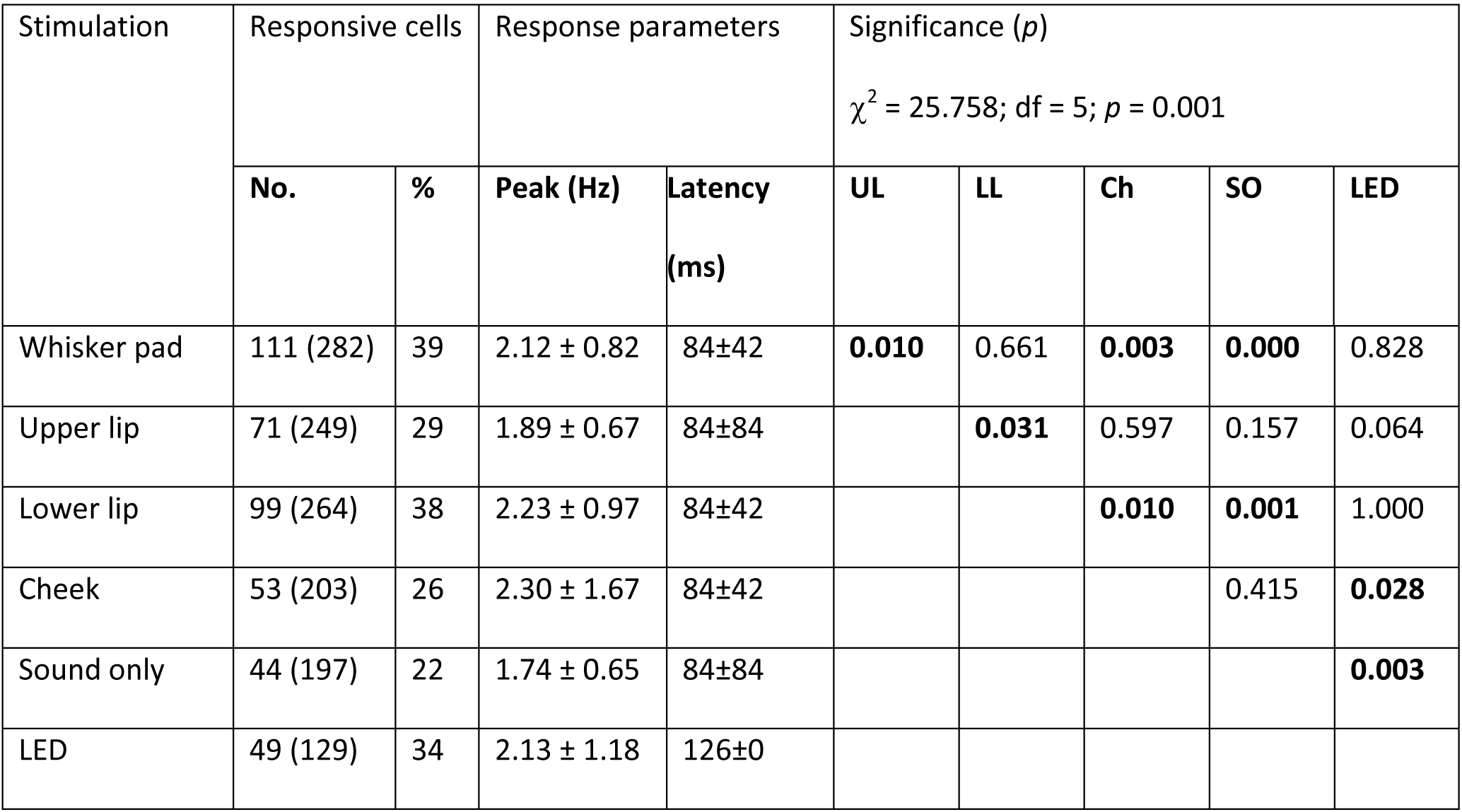
Purkinje cell responses to sensory stimulation in awake mice. Purkinje cells respond with complex firing to sensory stimulation (Fig. 4). For each type of stimulation, the number and percentage of statistically significantly responsive cells (peak response > average + 3 s.d. of baseline firing) is indicated (in brackets: total number of cells tested). The response peak and response latency are indicated as medians ± inter-quartile ranges. Some stimuli recruited more Purkinje cells with statistically significant responses than others. The differences in fractions of responsive Purkinje cells were statistically significant (6 × 2 χ^2^ test) and further tested using pair-wise Fisher’s exact tests (as the χ^2^ test was significant, no further correction for multiple comparisons was applied). Bold numbers indicate statistically significant values. UL = upper lip; LL = lower lip; Ch = cheek; SO = sound only.

### Convergence of sensory inputs

Next, we addressed the question whether Purkinje cells have a preference to respond to one or multiple types of stimulation. To this end, we made for each combination of two stimuli a scatter plot of the response strengths of each individual Purkinje cell. For each and every combination, correlation analysis revealed a positive correlation, implying that a stronger response to one stimulus typically implied also a stronger response to the other (Fig. 6A). These correlations, although weak, were statistically significant for all combinations except in two cases (i.e., upper lip vs. sound only and upper lip vs. visual stimulation; Table 2). The regression lines deviated from the 45° line, suggesting that, although there is a tendency at the population level to combine inputs, individual Purkinje cells can display specificity for a given stimulus. The Venn diagrams in Fig. 6B-C highlight the degree of overlap for all combinations of two and three stimuli. For all combinations we found some but no complete overlap. Remarkably, when comparing the observed overlap with the expected overlap based upon a random distribution, all combinations occurred more often than predicted. We performed a bootstrap analysis (see Methods) to infer statistical significance and indeed, most combinations were observed significantly more often than expected from a random distribution of inputs (Fig. 6B-C, Tables 3 and 4). This also included the visual stimulation, so that those Purkinje cells responding to a tactile stimulus typically were responsive to visual stimulation as well.

**Figure 6.**
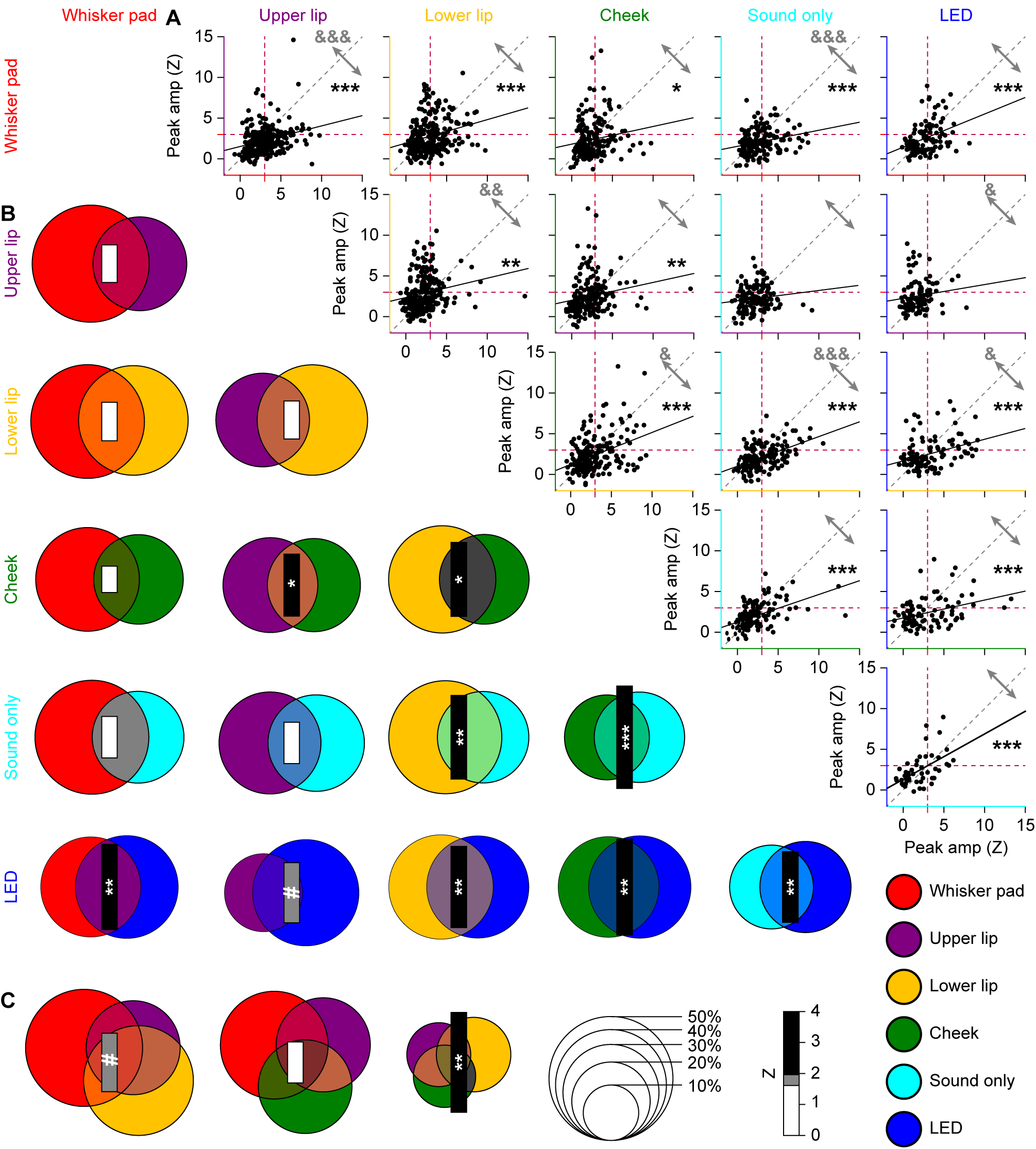
Convergence of sensory input on Purkinje cells. **(A)** In order to test whether sensory inputs converge on individual Purkinje cells in awake mice, we made pair-wise comparisons of the response amplitudes to two different stimuli per Purkinje cell (scatter plots). For all possible combinations, we found a positive slope of the linear regression analysis. For the majority of combinations, the correlation between response strengths was highly significant: * *p* < 0.05, ** *p* < 0.01 and *** *p* < 0.001 (Pearson correlation with Benjamini-Hochberg correction for multiple comparisons). Only upper lip vs. sound only and upper lip vs. visual stimulation were not significantly correlated. For this analysis, we included all Purkinje cells, whether they had a statistically significant response or not. The red dotted lines indicate a Z-score of 3, which we set as the threshold for significance (cf. Fig. 4B). They grey arrows indicate the fraction of observations above and below the unity line (grey dotted line). The relative strengths of each stimulus combination were compared in a pairwise fashion (Wilcoxon tests with Benjamini-Hochberg correction for multiple comparisons): & *p* < 0.05; && *p* < 0.01; &&& *p* < 0.001. **(B)** We performed a similar analysis focusing only on statistically significant responses (Venn diagrams). Again, all combinations had a positive Z score (as evaluated by a bootstrap method; see Methods), indicating more than expected convergence. The diameter of each circle indicates the fraction of Purkinje cells showing a significant response to that particular, colour coded stimulus. The size of the bar represents the Z score of the overlapping fraction. **(C)** The same for the combinations of three tactile stimuli. Overall, sensory streams tended to converge, rather than diverge, on Purkinje cells. # *p* < 0.10; * *p* < 0.05, ** *p* < 0.01 and *** *p* < 0.001 (Z test with Benjamini-Hochberg correction).

**Table 2.** Correlations between response probabilities. For each Purkinje cell, we made pair-wise comparisons of the Z scores of the amplitudes of the responses to different stimuli. For each pair of stimuli, the Pearson correlation coefficient (R) was calculated for the complete population of Purkinje cells subjected to that pair of stimuli. The *p* values of these correlations underwent Benjamini-Hochberg correction for multiple comparisons. These data are graphically represented in Fig. 5A. * *p* < 0.05; ** *p* < 0.01; *** *p* < 0.001

| Stimulation | Pearson correlation (R) |  |  |  |  |
| --- | --- | --- | --- | --- | --- |
|  | Upper lip | Lower lip | Cheek | Sound only | LED |
| **Whisker pad** | 0.272*** | 0.253*** | 0.166* | 0.257* | 0.358*** |
| **Upper lip** |  | 0.192** | 0.189** | 0.130 | 0.129 |
| **Lower lip** |  |  | 0.387*** | 0.513*** | 0.339*** |
| **Cheek** |  |  |  | 0.485*** | 0.362*** |
| **Sound only** |  |  |  |  | 0.523*** |

**Table 3.** Convergence of different sensory streams on individual Purkinje cells. For each type of stimulus, we compared the observed rate of convergence on individual Purkinje cells to chance level, using a bootstrap method based on the relative prevalence of responses to each stimulus (cf. **Fig. 5**). The difference from chance is indicated by *Z* values. All combinations occurred more often than expected (*Z* > 0). The *p* values of these correlations underwent Benjamini-Hochberg correction for multiple comparisons. * *p* < 0.05; ** *p* < 0.01

| Stimulation | Difference from chance (*Z*) |  |  |  |  |
| --- | --- | --- | --- | --- | --- |
|  | Upper lip | Lower lip | Cheek | Sound only | LED |
| Whisker pad | 1.21 | 1.25 | 0.85 | 1.34 | 2.75** |
| Upper lip |  | 1.23 | 2.00* | 1.34 | 1.94 |
| Lower lip |  |  | 2.39* | 2.70** | 2.62** |
| Cheek |  |  |  | 3.31** | 3.10** |
| Sound only |  |  |  |  | 2.28* |

**Table 4.** Convergence of different sensory streams on individual Purkinje cells. For each type of stimulus, we compared the observed rate of convergence on individual Purkinje cells to chance level, using a bootstrap method based on the relative prevalence of responses to each stimulus (cf. Fig. 5). The difference from chance is indicated by *Z* values. All combinations occurred more often than expected (*Z* > 0). The *p* values of these correlations underwent Benjamini-Hochberg correction for multiple comparisons. * *p* < 0.05; ** *p* < 0.01

| Stimulation | Difference from chance (Z) |
| --- | --- |
| **WP + UL + LL** | 1.89 |
| **WP + LL + Ch** | 1.32 |
| **UL + LL + Ch** | 3.22** |
| **WP + UL + LL + Ch** | 3.07** |

One might wonder whether Purkinje cells in crus 1 encode specific sensory events or, alternatively, respond indifferently to any external trigger. In this context, the response pattern to the sound only stimulus is especially noteworthy. A weak, but audible sound was generated by the piezo actuator used to deliver the tactile stimuli. Hence, all tactile stimuli also involved sound. Nevertheless, the Venn diagrams in Fig. 6B show that there are Purkinje cells that responded statistically significantly to sound only, but not to sound and touch delivered simultaneously. This could be taken as an argument against the input-specificity of Purkinje cells. However, one should keep in mind that the separation between “responsive” and “non-responsive” Purkinje cells is arbitrary (cf. Fig. 4B), and to be able to draw such a conclusion, there should be a consistent absence of preferred responses. The sound only stimulus was found to be the weakest stimulus (Figs. 4E and 5B) and pair-wise comparisons of the response strengths had a bias towards stimuli involving touch (Fig. 6A). This is further illustrated in a single experiment, directly comparing the responses evoked by whisker pad touch and sound only stimulation. Four randomly selected Purkinje cells from this experiment failed to show a response to sound only stimuli in the absence of touch responses (Fig. 7A-E). This was confirmed by the group-wise analysis of all Purkinje cells in this experiment, demonstrating a clear preference for the whisker pad stimulation over the sound only stimulus (Fig. 7F-G).

**Figure 7.**
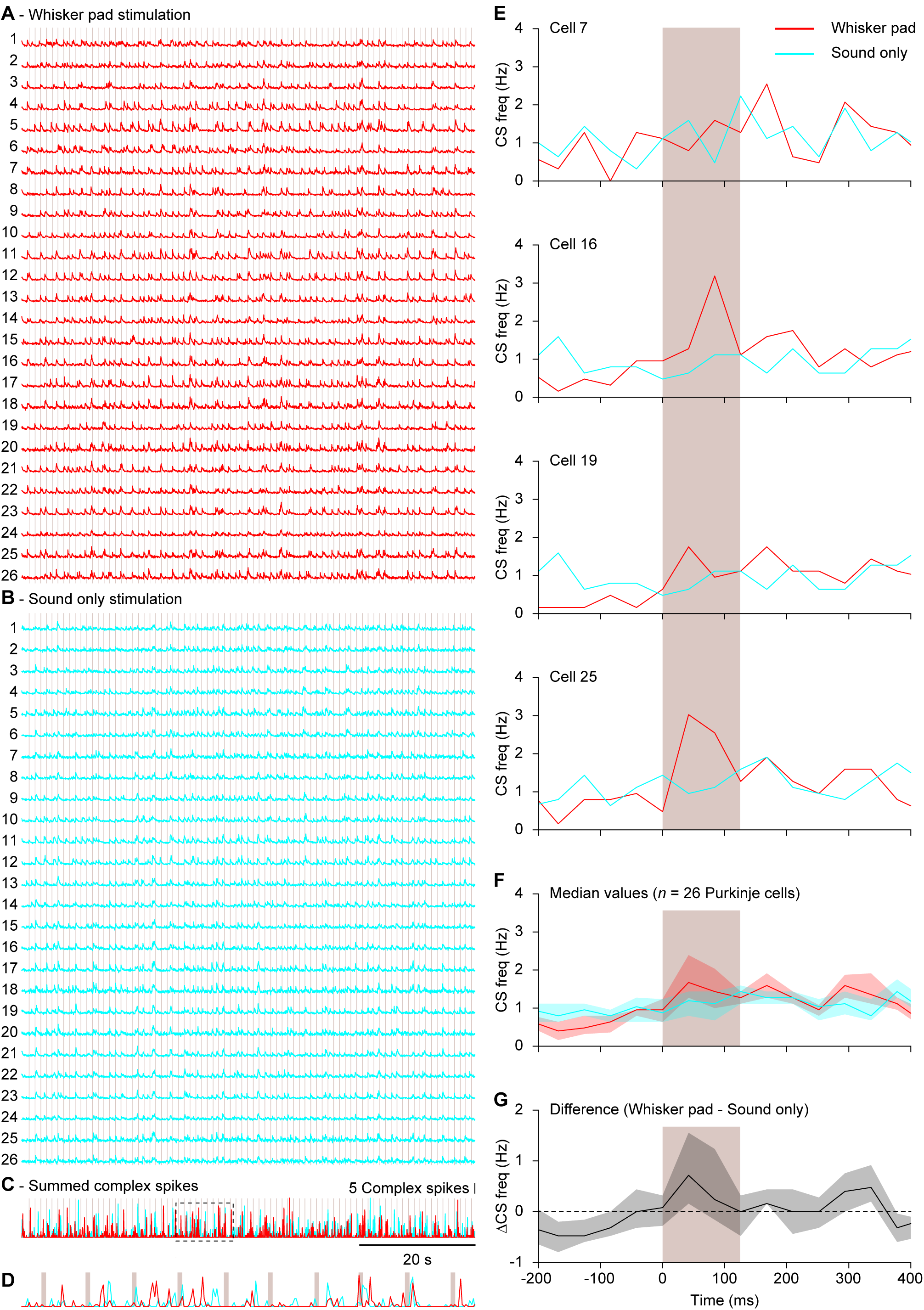
Sound only stimulation systematically recruited less complex spikes than tactile stimulation. Fluorescent traces of 26 Purkinje cells in a field of view during whisker pad **(A)** and sound only **(B)** stimulation. The moments of stimulation (at 1 Hz) are indicated by the vertical lines. Note that the sound only stimulation involved the sound of the mechanical device delivering tactile stimuli. Overall, whisker pad stimulation triggered more complex spike responses than sound only as illustrated by the sum of the events **(C)**. The boxed part (10 s) is enlarged in **D**. **(E)** Peri-stimulus time histograms (PSTHs) of four randomly selected Purkinje cells from the experiment illustrated in A and B. In each, the response to whisker pad stimulation was stronger than to sound only stimulation, which is also reflected in the median of the PSTHs of all 26 Purkinje cells **(F)** and the median difference between whisker pad and sound only stimulation **(G)**. The shaded areas indicate the inter-quartile range.

Thus, Purkinje cells in crus 1 seem to respond to multiple stimuli, although at the level of individual cells not all stimuli were equally effective in triggering complex spikes. To better disentangle the general responses from input-specific ones, we first focussed on the 188 Purkinje cells that received all four tactile stimuli. A full correlation analysis on this dataset (Fig. 8A) revealed similar results as the pair-wise comparisons shown in Fig. 6A. A partial correlation analysis found reduced correlations, confirming the existence of a common factor. However, some, but not all, pair-wise combinations occurred above chance level (Fig. 8B). If Purkinje cells respond to a generic sensory event, irrespective of stimulus location, such deviations from chance are not expected. This analysis, therefore, confirms the notion of a combination of a generic and an input-specific response. The impact of the common factor, representing a generic sensory trigger, was further investigated using principal component analysis. It turned out that the first component could explain 45% of the variance, which is significantly more than expected by chance (32%; Fig. 8C) and confirms that somatosensory input to Purkinje cells is only partially related to the specific stimulus location.

**Figure 8.**
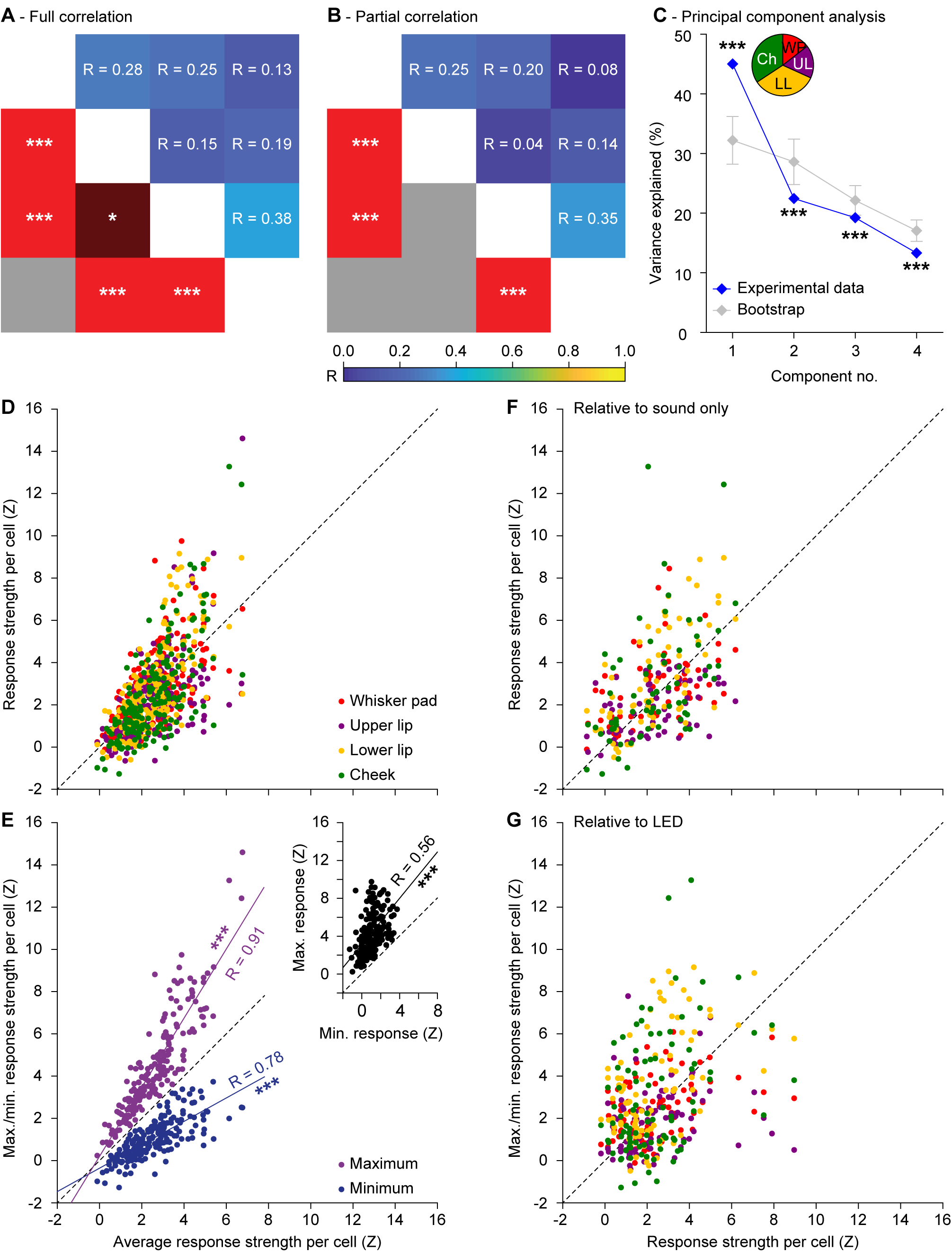
Purkinje cell excitation depends partially, but not completely, on generic sensory input. For the 188 Purkinje cells that received all four tactile stimuli, we calculated the full **(A)** and partial **(B)** correlation between the peak responses (in Z scores) of the four different tactile stimuli. This largely confirms the pair-wise correlations illustrated in Fig. 6A. Note, however, that the partial correlations are less pronounced than the full correlations, suggesting the existence of a common component reflecting general excitability, not specific for stimulus location. **(C)** Principal component analysis confirmed that a part of the observed variance can indeed be explained by a common factor, as the first principal component of the experimental data is significantly larger than that of bootstrapped data. Error bars indicate the 1-99% confidence interval. The inset shows the relative contributions of the different stimuli to the first principal component. The relatively weak stimuli (lower lip and cheek) were more in tune with the general excitability than the stronger stimuli (whisker pad and upper lip). **(D)** Scatter plot showing the correlation between average response strength to the four tactile stimuli vs. the four response strengths per Purkinje cell. The Purkinje cells on the left were insensitive to whatever tactile stimulus we presented, those on the right were sensitive to any tactile stimulus, but showed a bias towards one or a few stimulus locations. This bias becomes more obvious when plotting the minimum and maximum response per Purkinje cell **(E)**. Inset shows a strong correlation between the minimum and maximum response strength per Purkinje cell. R values come from Spearmańs correlation test. Similar plots were made comparing the sound only **(F)** and LED **(G)** stimulation vs. the four tactile stimuli. * *p* < 0.05, *** *p* < 0.001

To visualise the relation between a generic somatosensory response and input specificity, we made a scatter plot between the average peak responses to the four tactile stimuli versus the peak responses to each stimulus per Purkinje cell (Fig. 8D). Like in Fig. 4B, our data do not support unambiguous discrimination between “sensory” and “non-sensory” Purkinje cells. Several Purkinje cells did not show much of a tactile response to any given stimulus location, while there are also Purkinje cells that showed clear responses. The minimal and maximal responses diverge when the average response is stronger (Fig. 8E). As the average response depends on the minimal and maximal responses, this is not an independent measure. However, when correlating the minimal and maximal responses directly, a similar pattern emerges: stronger minimal responses correlate with even stronger maximal responses (Spearman’s correlation: R = 0.56, *p* < 0.001). In other words, Purkinje cells that respond to somatosensory stimulation at any given facial location are also prone to react to stimulation at another spot on the face. However, the Purkinje cells with a stronger “generic response” also had a stronger bias towards one or a few stimulus locations. Again, complex spike responses seem to be involved in the combination and not the segregation of sensory inputs, at the expense of a loss of input specificity. Similar analyses relating to auditory and visual stimulation showed that these conclusions generalize to other sensory modalities (Fig. 8F-G).

### Stimulus strength has limited impact on complex spike responses

To avoid recruiting responses from adjacent areas, we used weak stimulation strengths (Figs. 4 and 5). Under anaesthesia, the strength of a stimulus affects the complex spike response probability (Eccles *et al*., 1972; Bosman *et al*., 2010). This finding has been reproduced in awake mice using variations in duration and strength of peri-ocular air puff stimulation (Najafi *et al*., 2014). As a consequence, our approach using weak stimuli could have led to an underestimation of the number and spatial extent of sensory climbing fibre responses. To study whether the stimulus strength could also have affected our results, we performed an experiment in which we stimulated all whiskers mechanically with three different strengths, the largest of which was identical to the maximal stimulus we previously applied under anaesthesia (Bosman *et al*., 2010). The chosen stimulus intensities maximized the variation in kinetic energy, which was the most salient feature for barrel cortex neurons (Arabzadeh *et al*., 2004). Stimulus strengths were randomly intermingled (Fig. 9A-B). Raw traces indicate that, even at the ensemble level, weak stimuli do not trigger responses at every trial and spontaneous complex spike firing may occasionally appear as peaks prior to stimulus onset (Figs. 2C, 9C-D).

**Figure 9.**
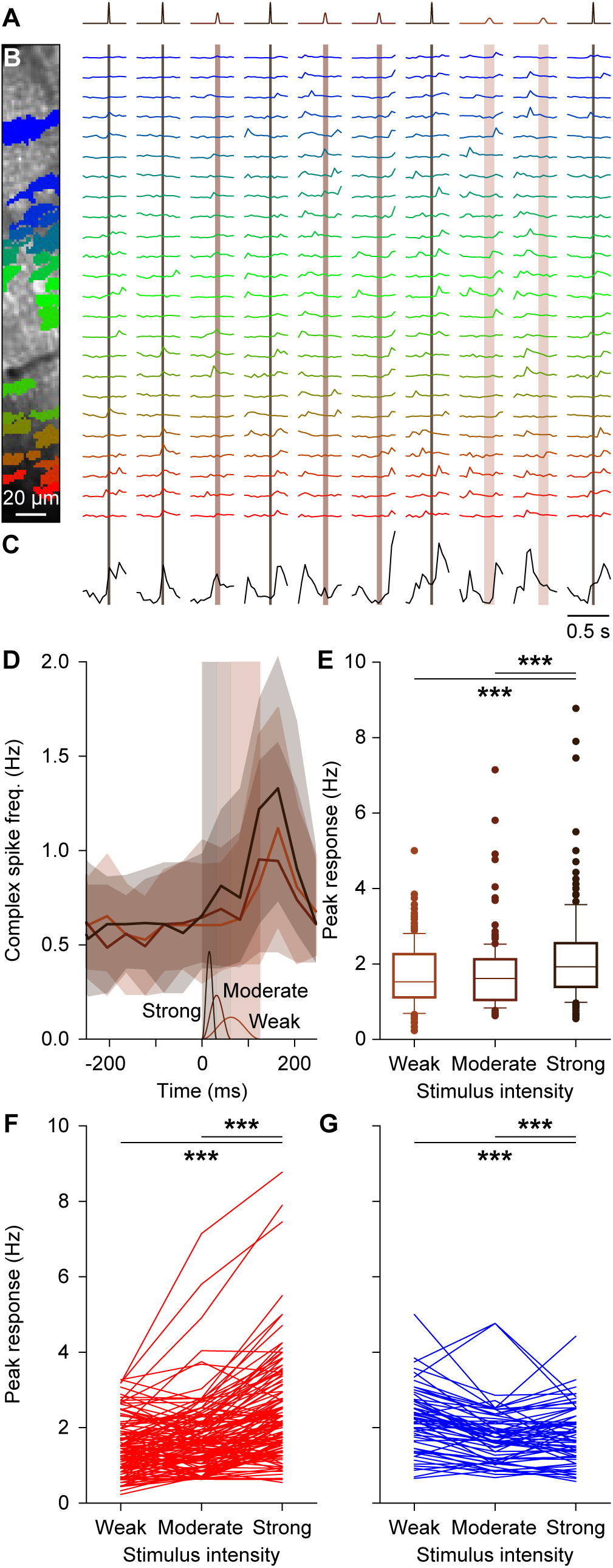
Stimulus strength has only a minor impact on complex spike responsiveness. **(A)** Movements of all large facial whiskers were performed using a piezo-actuator at three different speeds (weak: 1 mm displacement in 62 ms; moderate: 2 mm displacement in 31 ms; strong: 4 mm displacement in 16 ms). The stimulus sequence was randomly permuted. The recordings were made in awake mice. **(B)** Field of view with 24 identified and colour-coded Purkinje cells (left) and their corresponding fluorescent traces (right). Stimuli were presented every 2 s and in between trials the laser illumination was briefly blocked to avoid photobleaching. Note that the periods without laser illumination are not drawn to scale. The vertical shaded areas indicate stimulus duration (which was inverse with the stimulus strength). **(C)** Summed fluorescence trace composed of all 24 individual traces showing that not all trials evoked ensemble-wide responses. Some spontaneous, inter-trial activity was also observed. **(D)** The median number of complex spikes per frame (of 40 ms) per trial (shaded areas: inter-quartile ranges) for the three stimulus strengths show little difference for the weak and moderate stimulation. The time course and amplitude (1-4 mm) of the three stimuli is schematized at the bottom of the graph. Strong stimulation elicited about 30% more complex spikes, as evident from the peak responses for each stimulus intensity. **(E)** Box plots showing the response strength for the 209 significantly responsive Purkinje cells (out of 340 Purkinje cells that were measured in this way). **(F)** Response rates for all Purkinje cells that showed an increase in response rate with increased stimulus intensity. Only a few cells stand out in that they show a strong response that consistently increases with stimulus strength (lines on top). **(G)** The same for the Purkinje cells that showed a decrease in response strength with increasing stimulus intensity. *** *p* < 0.001 (Friedman’s test).

Of the 340 Purkinje cells tested in awake mice, 209 (61%) responded statistically significantly to at least one stimulus strength. In these 209 Purkinje cells, we compared the complex spike responses across three intensities. The weak and the moderate intensities (1 mm displacement reached in 62 ms and 2 mm displacement reached in 31 ms, respectively) showed a comparable number of responses and only the strong stimulus intensity (4 mm reached in 16 ms) evoked significantly more responses (*F*(2) = 57.160, *p* < 0.001, Friedman’s test) (Figs. 9D-E). Despite being 16 times faster (250 mm/s instead of 16 mm/s), the strongest stimulus recruited only 28% (measured as peak response) or 34% (measured as integral of whole response period) more complex spikes than the weakest stimulus (Fig. 9E-G). The same analysis on the whole population of Purkinje cells, including those that did not show a statistically significant response, revealed even less of an impact of stimulus strength (*data not shown*). We therefore conclude that the complex spike response encodes poorly the velocity of whisker displacement in awake mice and that – within boundaries – using stronger stimuli does not necessarily lead to qualitatively different results.

### Functionally equivalent Purkinje cells tend to group together

So far, we mainly found an abundance of weak and fairly non-specific complex spike responses to mild sensory stimulation. One way in which such weak responses at the cellular level could still have considerable effects at the network level would be when functionally equivalent Purkinje cells would lie together in microzones. Indeed, as the Purkinje cells of each microzone project to a group of adjacent neurons in the cerebellar nuclei (Voogd & Glickstein, 1998; Apps & Hawkes, 2009), population encoding of sensory events may be a form of functional signalling in the olivo-cerebellar system. Fig. 10 illustrates to what extent adjacent Purkinje cells have similar stimulus response probabilities. In this example, Purkinje cells responding strongly to whisker pad stimulation are grouped together, but such a spatial clustering does not seem to be perfect, as also a few strongly responsive cells are located in between less responsive Purkinje cells (Fig. 10A) and as upper lip-responsive Purkinje cells are sparsely distributed across the same area (Fig. 10B). We reasoned that spatial clustering should imply that two neighbouring cells have more similar response probabilities than randomly selected cells from the same field of view. This turned out to be the case, but only if we confined our focus to Purkinje cells with statistically significant responses (Z > 3) (Fig. 10C-D). Thus, especially the Purkinje cells with stronger responses to a given stimulus type were located closer than could be expected from a random distribution.

**Figure 10.**
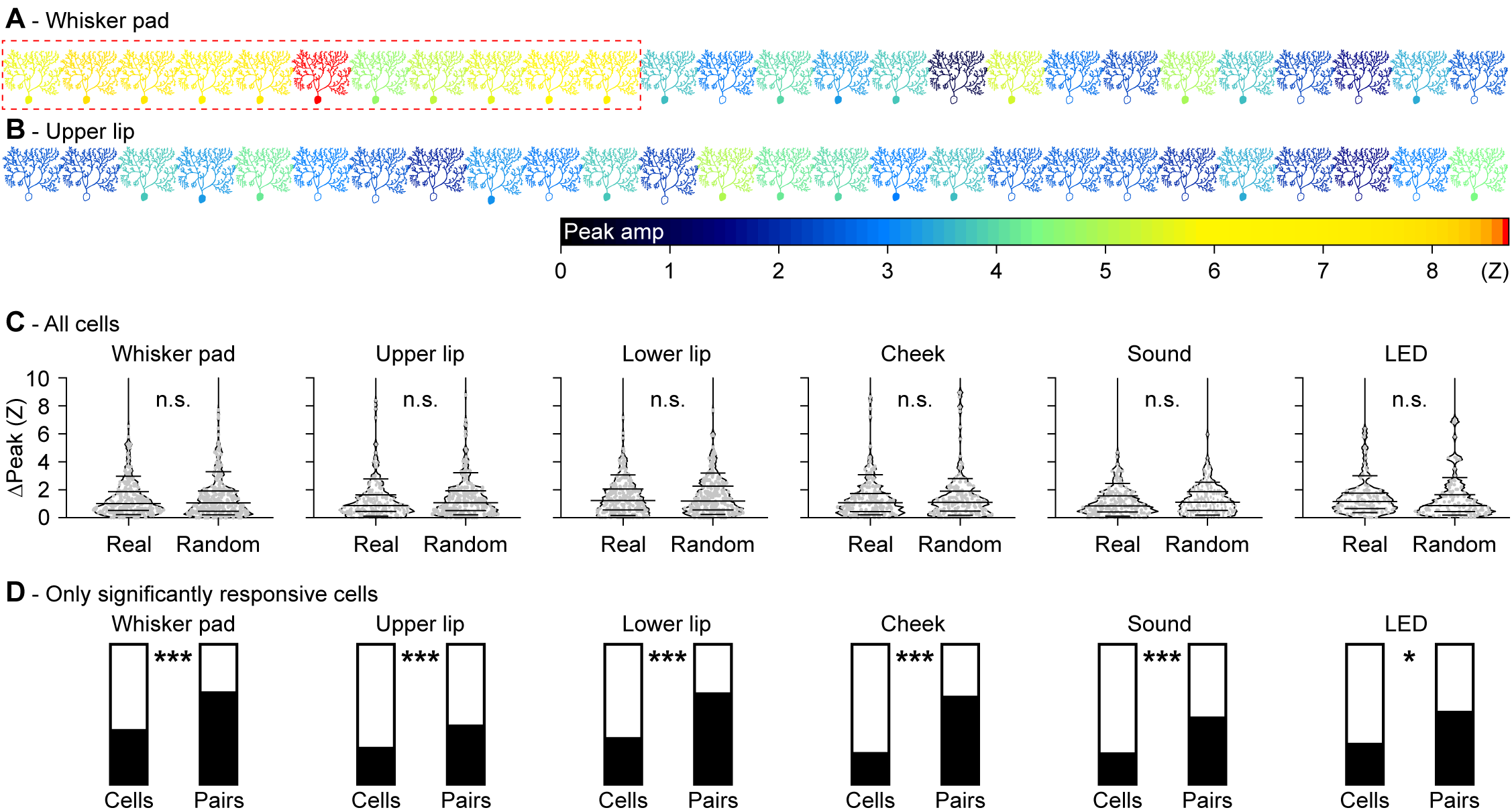
Purkinje cells encoding the same stimulus have a tendency to be spatially grouped. Schematic drawing of a field of view with 26 Purkinje cells organized in the medio-lateral direction of crus 1 in an awake mouse. The colour of each Purkinje cell corresponds to the maximal response to whisker pad **(A)** or upper lip **(B)** stimulation. Purkinje cells with a filled soma had a peak response with a Z score > 3 and were considered to be statistically significant, in contrast to those with an open soma. Responsive and non-responsive cells are generally intermingled, but a group of “strong responders” can be observed for whisker pad stimulation (red rectangle). **(C)** The anecdotal data in **A** suggest the presence of clusters of Purkinje cells encoding specific stimuli. For this to be the case, one would expect that neighbouring Purkinje cells have roughly similar response strengths. We found that this assumption does not hold as the differences in response strengths of neighbours could not be discriminated from randomly selected cells in the same recording if all Purkinje cells are considered (compared with bootstrap analysis based upon randomly chosen cell pairs within each field of view: all *p* > 0.8; Z test). Data are represented in violin plots, with the grey lines indicating the 10^th^, 25^th^, 50^th^, 75^th^ and 90^th^ percentiles. **(D)** When considering only the Purkinje cells with statistically significant responses, spatial grouping does occur. For each stimulus type, the black portion of the left bar indicates the fraction of Purkinje cells showing a significant response to that stimulus. The filled portion of the right bar indicates the fraction of the neighbours (always on the medial side) of these significantly responsive Purkinje cells that were also significantly responsive. As can be seen, this fraction is always substantially larger than the fraction of significantly responsive Purkinje cells, indicating a tendency of similar Purkinje cells to group together. Statistical significance was tested by comparing the fraction of Purkinje cells with statistically significant responses and the fraction of neighbours of Purkinje cells with statistically significant responses that showed statistically significant responses as well (after correction for border effects) using Fisher’s exact test and after Benjamini-Hochberg correction for multiple comparisons: * *p* < 0.05; *** *p* < 0.001.

### Single whisker responses in Purkinje cells

Using our tactile stimuli we demonstrated a tendency of nearby Purkinje cells to encode the same stimulus. To study whether this would hold true for even smaller receptive fields we turned to single whisker stimulation. Using single-unit electrophysiological recordings, we have previously shown that stimulation of a single whisker is sufficient to evoke complex spike responses (Bosman *et al*., 2010). Individual whiskers can be reliably identified across mice and as such they can be qualified as minimal reproducible receptive fields. We repeated our previous single-whisker stimulation experiments now using two-photon Ca^2+^ imaging focusing on five whiskers that were stimulated individually in a random sequence. To prevent spontaneous whisking and thereby interference by other whiskers, these experiments were (unlike all other experiments in this study) performed under anaesthesia. In general, the responses were specific, as many Purkinje cells responded to a particular whisker, but not to its neighbouring whiskers (Fig. 11A). Overall, of the 148 Purkinje cells tested, 31 (21%) responded significantly to only one of the five whiskers tested, 14 (10%) to two and 5 (3%) to three whiskers (Fig. 11B). Not all whiskers were equally effective in recruiting Purkinje cell responses: 23 cells (15%) responded significantly to C3 stimulation, but only 4 (3%) to C1 stimulation. Likewise, pairs of more anterior whiskers had higher chances to be encoded by the same Purkinje cell (Fig. 11C; Table 5). Thus, also for single whisker stimulation, there was a balance between sensory integration and specificity.

**Figure 11.**
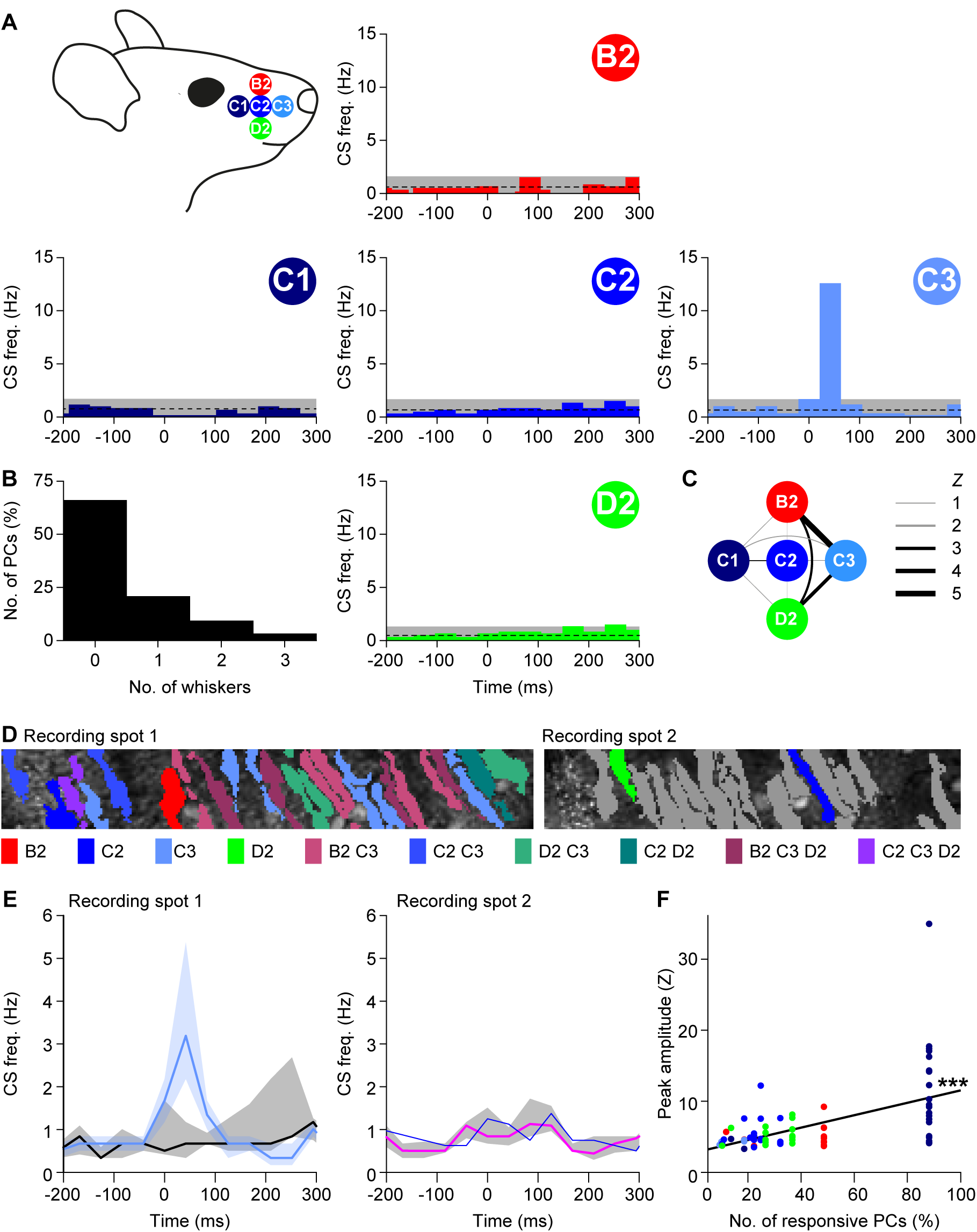
Purkinje cell responses to single whisker stimulation show weak clustering. **(A)** To investigate smaller receptive fields, we sequentially stimulated five of the large facial whiskers. To avoid interference with other whiskers during active movement, we performed these experiments under ketamine/xylazine anaesthesia. Most Purkinje cells, if responsive to single-whisker stimulation, responded only to one of the five whiskers **(B)**. This is illustrated by five peri-stimulus time histograms (PSTHs) from a single, representative Purkinje cell. This particular cell was sensitive to stimulation of the C3 whisker only. The average and 3 s.d. of the baseline firing are indicated (dashed line and grey area). **(C)** Purkinje cells that responded to more than one whisker were typically responsive to the more anterior whiskers (see also Table 5). The widths of the lines indicate the Z value of the occurrence of multiple responses per cell. **(D)** Two recording spots, in close proximity in crus 1 of the same animal, with the identified Purkinje cell dendritic trees. For each dendrite, the colour indicates the whisker(s) to which it was responsive (see legend below with grey denoting the absence of a statistically significant response). **(E)** For each of the two recording sites, the medians of the responsive and the non-responsive Purkinje cells are indicated (to the C3 whisker in the left panel and to the C2 whisker in the right panel). Note that only a single cell was responsive to C2 stimulation in recording spot 2. The shades indicate inter-quartile ranges. **(F)** Linear regression revealed that Purkinje cells that were surrounded by other Purkinje cells responsive to the same whisker (same colour code as in **A**) had a tendency to show stronger responses to stimulation of that whisker than Purkinje cells that were more isolated. The x-axis represents the fraction of Purkinje cells responsive to the particular whisker within the respective field of view. R = 0.521; *p* < 0.001.

**Table 5.** Purkinje cell responses to single-whisker stimulation in anaesthetized mice. Purkinje cells respond with complex firing to single-whisker stimulation (cf. **Fig. 10**). For each whisker, the number and percentage of responsive cells is indicated (in brackets: total number of cells tested). For each whisker, we compared the observed rate of convergence on individual Purkinje cells to chance level, using a bootstrap methods based on the relative prevalence of each stimulus. Indicated are the *Z* values. All combinations occurred more often than expected (*Z* > 0). Statistical significance is indicated after Benjamini-Hochberg correction for multiple comparisons. * *p* < 0.05; ** *p* < 0.01; *** *p* < 0.001

| Whisker | Responsive cells |  | Difference from chance (Z) |  |  |  |
| --- | --- | --- | --- | --- | --- | --- |
|  | No. | % | C1 | C2 | C3 | D2 |
| **B2** | 15 (148) | 10 | 0.630 | 0.227 | **5.130\*\*\*** | **2.052\*** |
| **C1** | 4 (148) | 3 |  | 0.803 | 0.782 | 0.630 |
| **C2** | 17 (148) | 11 |  |  | 0.222 | 0.208 |
| **C3** | 23 (148) | 16 |  |  |  | **3.100\*\*** |
| **D2** | 15 (148) | 10 |  |  |  |  |

Next, we examined the spatial distribution of responsive Purkinje cells. In Fig. 11D two nearby recording spots from the same mouse are shown. In recording spot 1, from which the example in Fig. 11A originates, all Purkinje cells responded to stimulation of at least one whisker. However, in the second recording spot, only two Purkinje cells responded: both to a single, but different whisker. For each recording spot we compared responsive vs. non-responsive Purkinje cells, taking the average + 3 s.d. of the baseline as threshold for responsiveness. This yielded a clear separation for the Purkinje cells in the first, but a rather poor one in the second recording spot (Fig. 11E). In terms of number of responsive Purkinje cells and response amplitudes, these two recordings, although made from the same lobule in the same animal, form relatively extreme examples in our dataset. When plotting the response strength versus the fraction of responsive Purkinje cells per field of view, we found a positive correlation (Pearson correlation: R = 0.52, *p* < 0.001; Fig. 11F), implying that Purkinje cells with stronger responses to a certain whisker tended to be surrounded by other Purkinje cells encoding the same whisker – in line with the much stronger responses in the first than in the second recording spot illustrated in Fig. 11D-E. Taken together, stimulating single whiskers revealed a similar organization as did the less specific stimuli (see Fig. 10) with a clear tendency of Purkinje cells with the same receptive field to be located close together. However, the spatial clustering was incomplete and especially weakly responsive Purkinje cells were found to be interspersed with completely unresponsive Purkinje cells.

### Functionally equivalent Purkinje cells fire coherently

In addition to spatial clustering, a second requirement for population encoding may be coherence of complex spike firing. While we refer to synchrony of two active Purkinje cells when they fire simultaneously within bins of 2 ms, we define coherence as firing simultaneously within bins of 40 ms. Time frames of 40 ms, which have historically also described as contemporaneous firing (Wylie *et al*., 1995), fall well within the subthreshold oscillation cycle of inferior olivary neurons *in vivo* (Khosrovani *et al*., 2007). Under anaesthesia, clusters of Purkinje cells tend to have increased coherence. These clusters organize in parasagittal stripes and their occurrence has been linked to zebrin bands (Sugihara *et al*., 2007; Ozden *et al*., 2009; Tsutsumi *et al*., 2015). To examine whether such parasagittal bands could also be detected in awake mice, we performed a set of imaging experiments with a larger field of view, but at a lower frame rate (15 Hz). Correlation analysis of these data confirmed the existence of parasagittally oriented clusters, although the demarcation of the clusters was not as sharp as observed under anaesthesia (cf. Ozden *et al*., 2009; Tsutsumi *et al*., 2015) (Fig. 12A-C).

**Figure 12.**
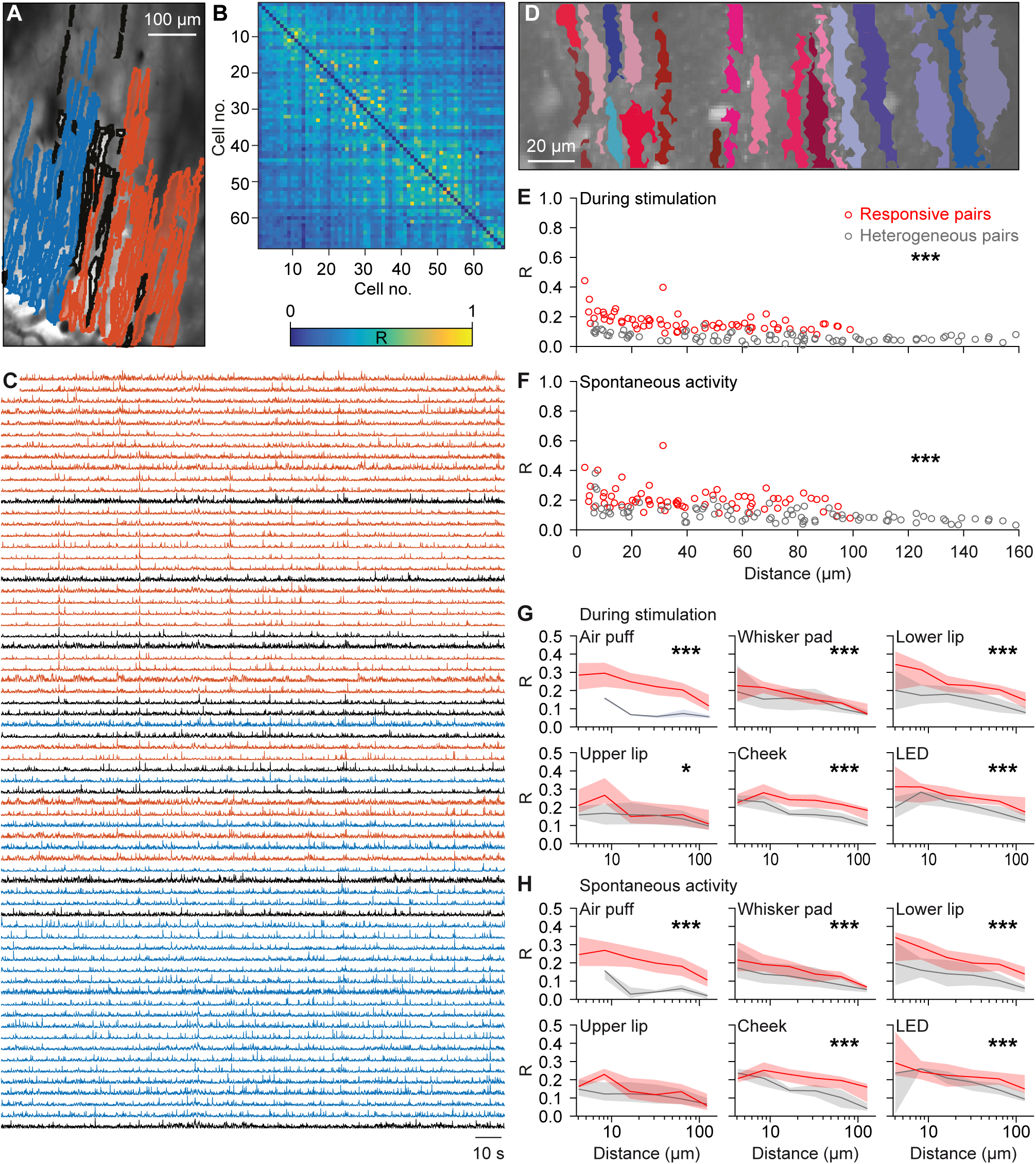
Purkinje cells encoding the same response are closer together. **(A)** To compare the extent of synchronous firing between the medio-lateral and the antero-posterior axes, we made recordings with a larger field of view. Identified Purkinje cell dendrites in a representative field of view colour-coded according to their membership of one of the two cluster identified with meta k means clustering (Dunn index = 0.85). The dendrites indicated in grey could not be contributed to either of the two groups. **(B)** Heat map of the pair-wise comparisons of the correlation between firing of the dendrites shown in A. Although two clusters were identified, it is clear that under our recording conditions, synchronous firing is not strictly related to a single micro-zone. **(C)** Raw traces of the neurons indicated in **A** and **B**. **(D)** As most of the variation was along the medio-lateral axis, we continued with smaller fields of view oriented along the medio-lateral axis. Representative field of view with segmented Purkinje cell dendrites. Non-responsive cells are depicted in shades of blue and responsive cells in shades of red during whisker pad stimulation. **(E)** For each pair of Purkinje cells we calculated the correlation coefficient (R) during 1 Hz whisker pad stimulation. The pairs of two Purkinje cells that were both responsive to whisker pad stimulation had on average a higher level of synchrony than the pairs connecting a responsive and a non-responsive Purkinje cell (*p* < 0.001; two-dimensional Kolmogorov-Smirnov test). The pairs consisting of two non-responsive Purkinje cells were excluded from this analysis. **(F)** Interestingly, even in the absence of sensory stimulation, the pairs of Purkinje cells that were both responsive to whisker pad stimulation maintained a higher level of synchrony than “heterogeneous pairs”. Thus, Purkinje cells with the same receptive field tended to fire more synchronously, even in the absence of stimulation. This analysis was expanded in the presence **(G)** and absence **(H)** of sensory stimulation for six different types of stimulation and illustrated as the median R value per distance category (six bin values of equal distance at a log scale). The shaded areas represent the inter-quartile ranges. * *p* < 0.05; *** *p* < 0.001 (Kolmogorov-Smirnov tests)

As most of the variation could be found along the medio-lateral axis, we proceeded with smaller field of views along the medio-lateral axis to allow for a higher temporal resolution. We applied coherence analysis on spontaneous and modulation data obtained from Purkinje cells with significant responses in awake mice. Pairs of Purkinje cells with significant responses upon whisker pad stimulation showed a significantly increased level of coherence (*p* < 0.001; two-dimensional Kolmogorov-Smirnov test) compared to that of heterogeneous pairs (i.e. pairs of one responsive and one non-responsive Purkinje cell) (Fig. 12D-E). When we examined the firing pattern of the same pairs in the absence of sensory stimulation, we found similar results (Fig, 12F), indicating that it is not the sensory input per se that directs coherence. These findings were confirmed in the whole population, taking also the other tactile and visual stimuli into account (Fig. 12G-H). Only stimulation of the upper lip, the area least represented among the recorded Purkinje cells (Fig. 5B), revealed less of a discrimination between significantly responsive and heterogeneous pairs. Hence, we conclude that not only spatial clustering but also coherence patterning of functionally equivalent Purkinje cells may facilitate population encoding.

### Population responses

In general, the level of coherence among functionally equivalent Purkinje cells was relatively low, with a correlation coefficient seldom exceeding 0.3 (Fig. 12). However, in this analysis we did not discriminate between complex spike firing in response to sensory stimulation and complex spikes occurring spontaneously during inter-trial intervals. Therefore, we subsequently quantified the distribution of complex spikes over time, segregating both types of spikes. This is illustrated for a representative field of view in Fig. 13A. For each frame, we summed all complex spikes of the 17 Purkinje cells in this field of view and subsequently made an aggregate peri-stimulus time histogram of all these Purkinje cells, whereby we colour-coded the number of complex spike recorded per frame (see also Romano *et al*. {Romano, 2018 #4219}). The darker the colour, the more complex spikes occurred simultaneously. Clearly, the darker colours – and thus the stronger coherence – were observed during the stimulus response period. This occurrence of coherent firing over time was compared with a random redistribution of the spikes per Purkinje cell (based upon a Poisson distribution). Note that an equal distribution would imply on average less than two complex spikes being fired during each frame of 40 ms, indicating the highly patterned distribution found during the experiments. The grey bars in Fig. 13B indicate the level of coherence that could be expected by chance, while the red ones indicate highly unlikely values. This shows that especially the higher levels of coherence are task-related, while spatially isolated firing occurs irrespective of stimulation.

**Figure 13.**
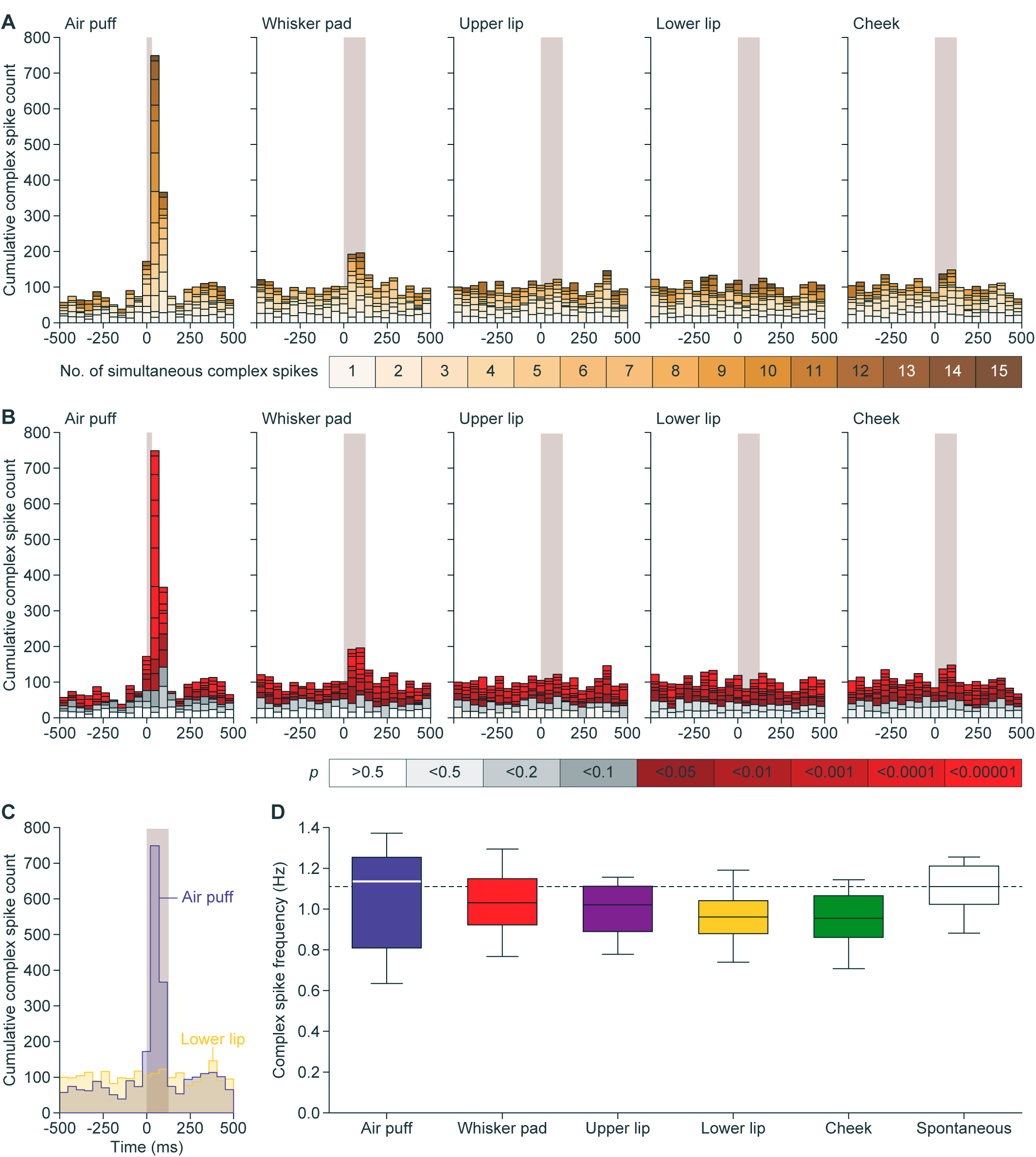
Purkinje cells encode strong and weak sensory stimulation via synchronous firing. **(A)** Aggregate peri-stimulus time histograms (PSTHs) show that coherent firing of complex spikes predominantly occurs following sensory stimulation. For each field of view, we calculated the number of complex spikes occurring per frame, summing those of all Purkinje cells in that field of view. Subsequently, we made aggregate PSTHs where the colour of each bin refers to the number of dendrites simultaneously active. In this field of view, 17 Purkinje cells were measured. Of these, 17 (100%) reacted to air puff, 12 (71%) to whisker pad, 3 (18%) to upper lip, 4 (24%) to lower lip and 1 (6%) to cheek stimulation. **(B)** Based upon a Poisson distribution of complex spikes over all dendrites and bins, one would expect between 0 and 3 simultaneously active dendrites (grey bars). The red bars indicate events involving more dendrites simultaneously than expected from a random distribution. Thus, the sparse firing as expected by chance is relatively constant throughout the trials, but the simultaneous activity of multiple dendrites is strongly enhanced following sensory stimulation. **(C)** A direct overlay of the aggregate PSTHs in response to air puff and lower lip stimulation shows that the strong response found after air puff stimulation comes at the expense of intertrial complex spikes (152 trials per condition). **(D)** For equally long recordings in the presence of different types of stimulation, equal complex spike frequencies were observed as during spontaneous activity (F(2.544, 20.348) = 2.561, *p* = 0.091, repeated measures ANOVA), indicating that sensory stimulation results in a temporal re-ordering of complex spikes, rather than to the production of more complex spikes.

Examination of the aggregate PSTHs confirms what could already be seen in Fig. 4D, in that there is a trend of reduced inter-trial firing for those stimuli with a relatively strong response (Fig. 13A-B). In this experiment, we plotted the responses to air puff stimulation, which recruited statistically significant responses in 17 out of 17 Purkinje cells, on top of those to lower lip stimulation, which recruited only 1 of the 17 Purkinje cells, illustrating the reduced baseline firing during air puff stimulation (Fig. 13C). When we calculated the average complex spike firing for each stimulus condition over the whole population of recorded Purkinje cells, we found that to be remarkably constant. Even 1 Hz air puff stimulation, able to recruit strong complex spike responses, did not result in increased complex spike firing as compared to an epoch without any form of stimulation (Fig. 13D), pointing towards a homeostatic mechanism within the inferior olive that balances out complex spike firing over longer time intervals.

### Homeostasis of complex spike firing

To further study the impact of complex spike homeostasis, we averaged the PSTHs of all Purkinje cells with statistically significantly responses to air puff stimulation and compared these to the firing rate in recordings made in the absence of stimulation (using pseudo-triggers generated at the same 1 Hz rate). This pair-wise comparison confirmed that the increase in complex spike firing during the sensory-induced responses comes at the expense of inter-trial firing. The same was true for the milder whisker pad stimulation. However, because the responses were weaker, the effect on the inter-trial firing was less than that following strong air puff stimulation. The homeostatic effect was also observed following visual stimulation, although this type of stimulation induced an oscillatory response, making the effect less visible (Fig. 14A-B). Taking the whole population into account, thus also the Purkinje cells without a statistically significant response, there proved to be a correlation between the peak of the stimulus response and the decrease in inter-trial firing (R = 0.40, *p* = 0.001 and R = 0.31, *p* = 0.001, Pearson correlations for air puff and whisker pad stimulation, respectively; Fig. 14C). Only following visual stimulation, this correlation was less obvious (R = 0.20, *p* = 0.201, Pearson correlation after Benjamini-Hochberg correction), possibly due to the oscillatory responses evoked by visual stimulation.

**Figure 14.**
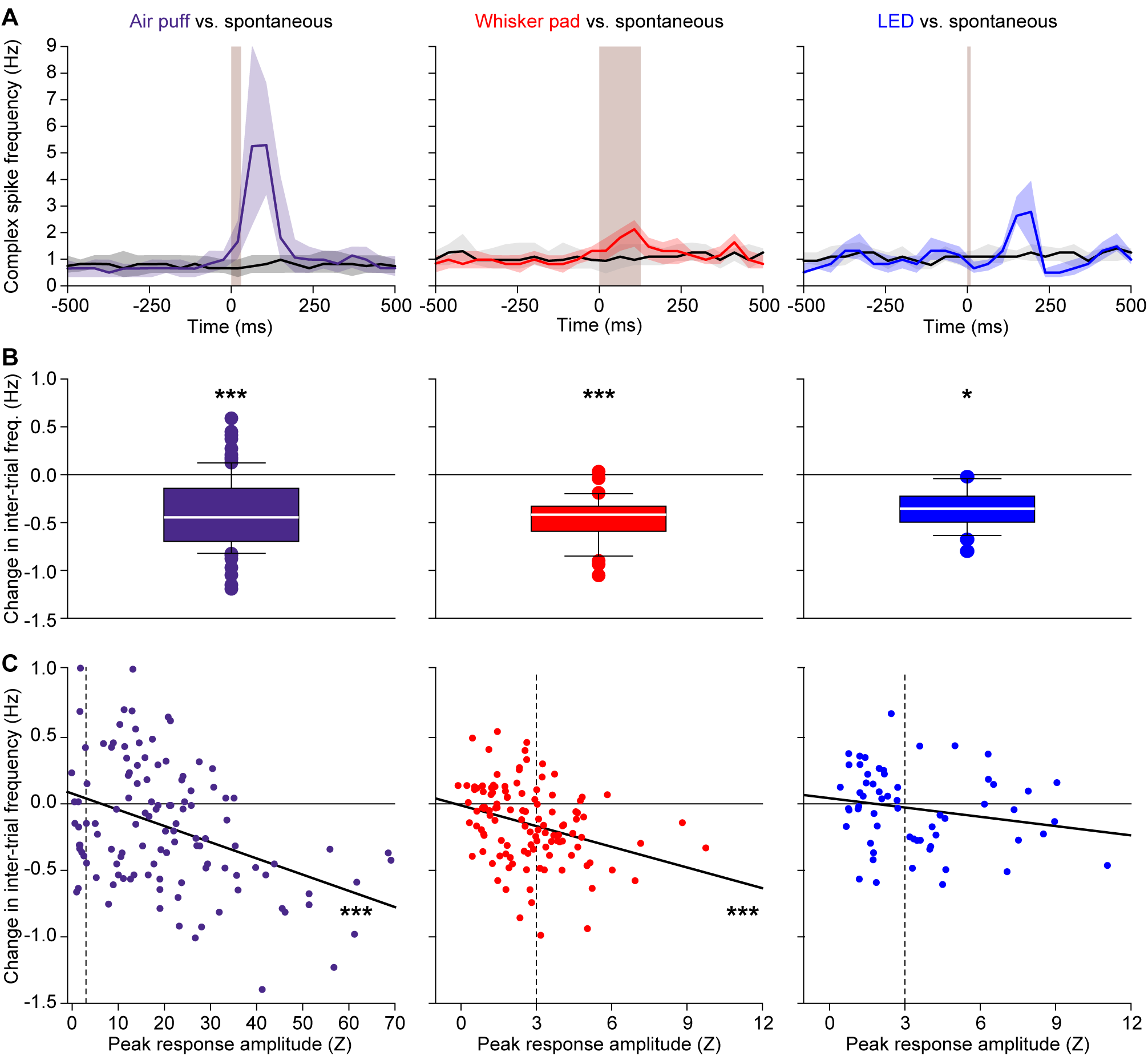
Sensory stimulation results in a temporal re-ordering of complex spikes. **(A)** The temporal distribution of complex spikes was compared in a pairwise fashion between sessions with sensory stimulation and sessions without. For this analysis, we included only Purkinje cells that displayed a statistically significant response to the stimulus involved (*n* = 102 for air puff, *n* = 45 for whisker pad and *n* = 27 for visual stimulation). The spontaneous recordings were analyzed by creating *post hoc* pseudo-stimuli at the same 1 Hz frequency as during sensory stimulation. Shown are the medians of the peri-stimulus time histograms. The shaded areas indicate the inter-quartile ranges. **(B)** The reduction in baseline firing, measured during the −500 to −250 ms interval, was significant in all cases (Wilcoxon matched-pairs test after Benjamini-Hochberg correction for multiple comparisons). **(C)** The larger the response amplitude, the stronger the reduction in inter-trial firing (Pearson correlation tests after Benjamini-Hochberg correction for multiple comparisons). This analysis was performed on all Purkinje cells (*n* = 117 for air puff and whisker pad stimulation and *n* = 60 for LED stimulation; dotted lines mark the criterion for statistical significance at Z = 3). Note the differences in the x-axis scaling with the air puff evoking relatively stronger responses. * *p* < 0.05; *** *p* ≤ 0.001

## Discussion

Complex spike firing may appear notoriously unpredictable as its spontaneous frequency is low, yet sustained, and its response rate is at best moderate, with large jitters and an ambiguous relation to stimulus strength. This raises the question as to how the inferior olive relays its signals over time. Possibly, simultaneous complex spike firing by neighbouring Purkinje cells might jointly represent a stimulus, together covering the required signalling for a particular temporal domain (Sasaki *et al*., 1989; Lang *et al*., 1999; Sugihara *et al*., 2007; Ozden *et al*., 2009; Schultz *et al*., 2009). Furthermore, the spatial relation between somatotopic patches and parasagittally oriented microzones has not been clarified in terms of complex spike signalling, and it is not understood to what extent this relation also depends on the temporal context in which the signals are generated.

Here, we investigated at the level of individual Purkinje cells whether encoding of somatosensory input from different facial areas in lobule crus 1 occurs in striped microzones or instead follows a more fractured arrangement. Mild touches at localized facial areas revealed a loose version of fractured somatotopy. Purkinje cells with the same receptive field tended to be located in each other’s neighbourhood, but the spatial organization was not very strict, as highly responsive Purkinje cells were sometimes observed amidst non-responsive ones. Functionally equivalent, adjacent Purkinje cells showed increased coherence in their complex spike firing, in particular in response to sensory stimulation. Moreover, homeostatic mechanisms were engaged, ensuring that over longer periods complex spike firing rates are constant, making the short-lived coherent responses more salient.

### The functional role of climbing fibre activity

Clinical manifestations of inferior olivary dysfunction range from ataxia and tremor to autism spectrum disorders (Llinás *et al*., 1975; Samuel *et al*., 2004; Bauman & Kemper, 2005; Welsh *et al*., 2005; Lim & Lim, 2009; De Gruijl *et al*., 2013). Climbing fibre-evoked complex spikes are also essential for the proper timing, size and direction of movements as well as for the encoding of expected and unexpected deviations from planned movements (Wang *et al*., 1987; Simpson *et al*., 1996; Kitazawa *et al*., 1998; Ito, 2013; Yang & Lisberger, 2014; Herzfeld *et al*., 2018; Romano *et al*., 2018). In addition, climbing fibre activity is crucial for cerebellar learning by controlling synaptic plasticity at a wide variety of synapses in the molecular layer of the cerebellar cortex (Ito & Kano, 1982; Ito, 2003; Coesmans *et al*., 2004; Van Der Giessen *et al*., 2008; Gao *et al*., 2012). Despite these many functions, complex spike firing is remarkably stable over longer intervals, suggesting that the impact of complex spikes is strongly context-dependent.

This context is provided by the synaptic inputs to the inferior olive that relay information from excitatory ascending and descending pathways as well as inhibitory projections from the hindbrain (De Zeeuw *et al*., 1998). Both types converge on each individual spine present on the dendrites of the inferior olivary neurons (De Zeeuw *et al*., 1989; De Zeeuw *et al*., 1990). The inhibitory input predominantly originates from the cerebellar nuclei, implying that part of the context is mediated by one of the target regions of the climbing fibres themselves. This feedback is engaged in a closed loop, as individual climbing fibres of each olivary subnucleus project to the Purkinje cells located within a specific parasagittal zone (Sugihara *et al*., 2001) that converge on the neurons in the cerebellar nuclei that project back to the same olivary subnucleus where the loop started (Groenewegen *et al*., 1979; Voogd & Glickstein, 1998; Apps *et al*., 2018). Each module can be further subdivided into microzones within which complex spike coherence is clearly enhanced upon strong stimulation (Ozden *et al*., 2009; Tsutsumi *et al*., 2015), a phenomenon that is promoted by strong electrotonic coupling within olivary glomeruli (Sotelo *et al*., 1974; Ruigrok *et al*., 1990; Devor & Yarom, 2002; De Gruijl *et al*., 2014). Under anaesthesia, complex spike coherence adheres relatively well to the delineation of zebrin bands (Sugihara *et al*., 2007; Ozden *et al*., 2009; Tsutsumi *et al*., 2015). In awake mice, the spatial and functional demarcation between adjacent coherent clusters of Purkinje cells is diminished (Fig. 12B and De Gruijl *et al*. (2014)), probably due to the more diverse pattern of synaptic input to the inferior olive present in awake mice. Thus, although anatomical and functional data support the microzonal organization, the activity patterns are best understood as produced dynamically depending on the behavioural context that is transmitted via cerebellar and extra-cerebellar synaptic input (Negrello *et al*., 2018).

A particularly interesting model system to study differential functional roles of climbing fibre activity is classical eyeblink conditioning. Subjects can readily learn to associate a previously neutral stimulus with an aversive puff to the eye. Before training, complex spikes virtually only occur after the unconditioned air puff stimulus. However, after training, the initial neutral stimulus, the conditioned stimulus, also reliably triggers complex spike responses at the onset time of the conditioned response (Halverson *et al*., 2015; Ohmae & Medina, 2015; Ten Brinke *et al*., 2015). It appears that the nature of the conditioned stimulus, whether auditory, visual or tactile (Hilgard & Marquis, 1936; McCormick & Thompson, 1984; Das *et al*., 2001; Galvez *et al*., 2006) is of subordinate relevance in this respect and that the main goal of these conditioned stimulus related complex spikes is to further improve the conditioned motor response (Ohmae & Medina, 2015; Ten Brinke *et al*., 2015; Ten Brinke *et al*., 2019). However, the conditioned stimulus does not generalize: subjects trained with a visual conditioned stimulus will not blink after receiving an auditory stimulus for instance, although the associative learning process for a second stimulus goes faster than for the first (Kehoe & Holt, 1984; Campolattaro & Freeman, 2009; Campolattaro *et al*., 2015). Such a flexible arrangement would be very much in line with our data. In a naïve mouse, multiple weak sensory streams as well as internally generated sensorimotor prediction signals may converge on individual Purkinje cells (present study; Heffley *et al*. (2018)). Consequently, most sensory stimuli trigger only weak complex spike responses. Given the behavioural saliency, one or a few pathways may grow stronger and account for the Purkinje cells with a strong complex spike response. Thus, there is a balance between generalized input, encoding basically any sensory or internally generated event, and input-specificity.

To what extent motor related enhancement of complex spikes responses as observed during eyeblink conditioning are module-dependent remains to be shown. In principle such responses might be dominant in zebrin-negative modules, in which simple spike responses are suppressed during learning (De Zeeuw & Ten Brinke, 2015) and in which climbing fibre responses can further facilitate such suppression (Ten Brinke *et al*., 2015). In zebrin-positive zones, in which the expression of learned responses are more driven by increases of simple spike responses (De Zeeuw & Ten Brinke, 2015; Voges *et al*., 2017; Romano *et al*., 2018), the opposite might occur. For example, it has been shown that olivary activity can be inhibited and gated when sensory stimuli are applied in a predictable fashion during self-generated locomotion (Gibson *et al*., 2004).

### Receptive fields of Purkinje cells and population coding

Several existing maps of somatosensory representations of complex spike activity suggest a fractured somatotopy (Miles & Wiesendanger, 1975; Rushmer *et al*., 1980; Castelfranco *et al*., 1994). These maps were typically created by establishing, for each recording position, the strongest input region, disregarding information on convergence of multiple inputs. These studies made clear, however, that climbing fibres could have receptive fields of widely different sizes, expanding to structures as large as a whole limb (Thach, 1967; Leicht *et al*., 1973). Here, we show at the single cell resolution that climbing fibres indeed do convey somatosensory input from different areas, with Purkinje cells having the same receptive fields being located preferably in each other’s proximity.

Our data reveal that classification of Purkinje cells into responsive and non-responsive cells is at best disputable, with the large majority of Purkinje cells typically showing relatively weak responses (Fig. 4B). Whisker pad stimulation elicits the strongest responses in lateral crus 1 (Romano *et al*., 2018), while most of the current data were collected in more medial parts of crus 1. Consequently, only a few Purkinje cells included in the present study showed a strong response to whisker stimulation. Remarkably, these were the ones that did depend on stimulus strength, in contrast to the majority of the weakly responsive Purkinje cells (Fig. 9F). This may point to the existence of two groups of Purkinje cells: some that show complex spike responses that clearly depend on stimulus strength, in line with previous reports (Eccles *et al*., 1972; Bosman *et al*., 2010; Najafi *et al*., 2014), and others that display only a weak to moderate sensitivity to a specific sensory input, independent of stimulus strength.

For the Purkinje cells with strong responses, a function in the timing and/or fine-tuning of motor responses is likely. Complex spikes have been shown to adjust saccadic eye movements on a trial-by-trial basis (Yang & Lisberger, 2014; Herzfeld *et al*., 2018). They are also correlated to the amplitude of reflexive whisker movements (Romano *et al*., 2018). The abundance of weak responses could, as discussed above for zebrin-negative modules, possibly be explained in terms of forming a substrate for cerebellar learning, where specific training would strengthen particular pathways as, for instance, occurs during classical eyeblink conditioning (Ten Brinke *et al*., 2015).

An alternative, but not mutually exclusive, explanation for the presence of weak responses could be multi-sensory integration. Analysing the response rates of Purkinje cell ensembles revealed that the density of responsive cells is crucial for shaping the population response. We propose that the weak spatial clustering of Purkinje cells encoding the same stimulus, being interspersed with Purkinje cells receiving input from other sources, and the tendency of coherent firing of Purkinje cells encoding the same stimulus both contribute to the creation of a heterogeneous map of Purkinje cells, where each area encodes a particular functional set of inputs. Each of the properties of the complex spikes seems rather insignificant in isolation, but in combination may result in specific and robust population encoding.

### Non-tactile inputs

Light or sound can act as conditional stimulus to recruit increasingly more climbing fibre activity during learning, highlighting the flexibility of this pathway (Ohmae & Medina, 2015; Ten Brinke *et al*., 2015). In naïve animals, auditory and visual input have been shown to project predominantly to the vermis (Snider & Stowell, 1944), but we demonstrate here that visual and auditory input can converge on the same Purkinje cells that encode somatosensory input in crus 1 – also in naïve mice – further strengthening the notion that climbing fibre activity can act as an integrator of contextual input. With visual stimulation the latency was significantly longer than expected. Probably, the presented visual stimulus is not directly relayed from the retina, but a descending input from the visual cortex or other higher brain regions such as the mesodiencephalic junction (De Zeeuw *et al*., 1998).

### Homeostasis of complex spike frequency

The overall firing rate of complex spikes was stable across conditions, with response peaks being compensated by reduced inter-trial firing (see also Negrello *et al*. (2018)). The stronger the response peak, the less inter-trial firing, leading to homeostasis of complex spike firing over longer time periods. Intriguingly, the complex spikes during the inter-trial intervals were predominantly fired by a few, dispersed Purkinje cells and the response peak was largely due to increased coherence. It seems therefore that Purkinje cells display a basic complex spike firing rate, which in the absence of functional behaviour is not coherent with adjacent Purkinje cells. In view of the strong impact of complex spikes on synaptic plasticity (Ito & Kano, 1982; Ito, 2003; Coesmans *et al*., 2004; Gao *et al*., 2012), this could subserve homeostatic functions. Salient stimuli are largely encoded by coherent firing, making them different from non-stimulus related activity.

Short-lived excursions from the stereotypic 1 Hz complex spike frequency occur often.Many studies have documented a 10 Hz rhythm in complex spike firing (Bell & Kawasaki, 1972; Wylie *et al*., 1995; Lang *et al*., 1999; Blenkinsop & Lang, 2006), which is sustained over longer periods after harmaline application (Llinás & Volkind, 1973). Also our current data (e.g., see double spike in Fig. 1D) show that the inferior olive as well as the Purkinje cells are physically able to fire much faster than at 1 Hz. Several mechanisms may contribute to the homeostatic control of inferior olivary spiking, including the previously mentioned olivo-cerebellar loops. Disrupted Purkinje cell activity can affect climbing fibre activity (Chen *et al*., 2010) and reduced inferior olivary activity leads to enhanced simple spike activity (Montarolo *et al*., 1982), which in turn dampens activity in the GABAergic neurons of the cerebellar nuclei that controls the inferior olive (Chaumont *et al*., 2013; Witter *et al*., 2013). The interplay between glutamatergic input to the neurons of the inferior olive, intracellular signalling and electrotonic coupling may also lead to homeostatic control of complex spike firing via PKA and ßCaMKII (Mathy *et al*., 2014; Bazzigaluppi *et al*., 2017). Overall, the inferior olive functions at the cross road of well-defined and rigid anatomical structures and highly dynamic synaptic input, while subject to homeostatic control, the sources of which still are partly to be determined.

## Additional information

### Author contributions

All experiments were performed at the Department of Neuroscience of the Erasmus MC, Rotterdam, The Netherlands. The experiments were designed by C.J., L.W.J.B., T.M.H., P.M., M.N. and C.I.D.Z., performed by C.J. and P.M. and analysed by all authors. The manuscript was written by C.J., L.W.J.B., T.M.H., M.N. and C.I.D.Z. with contributions from all authors. All authors have approved the final version of the manuscript and ensure the accuracy and integrity of the data presented. All persons with a significant contribution to this study are listed as authors.

## Acknowledgements

The authors wish to thank A.C.H.G. IJpelaar for technical assistance and Dr. M. Schonewille for help with recording ocular movements during stimulation.

